# Dynamics of the epigenetic landscape during development and in response to drought stress in sorghum

**DOI:** 10.1101/2023.04.24.537601

**Authors:** Yongfeng Hu, Chao He, Yuning Shen, Gongjian Zeng, Siteng Bi, Quanjun Huang, Xiner Qin, Zhuying Deng, Zhengquan He, Xiangling Shen

**Affiliations:** Key Laboratory of Three Gorges Regional Plant Genetics & Germplasm Enhancement (CTGU)/ Biotechnology Research Center, College of Biological and Pharmaceutical Sciences, China Three Gorges University, Yichang, Hubei 443002, China; National Key Laboratory of Crop Genetic Improvement, Hubei Hongshan Laboratory, Huazhong Agricultural University, Wuhan, China

**Keywords:** sorghum, histone modification, H2A.Z, root and leaf, drought stress

## Abstract

*Sorghum bicolor* is a C4 plant with the characteristics of high stress tolerance, which may be conferred partly by the underlying epigenetic mechanism unique to sorghum. In this study, we revealed some epigenomic features in sorghum that have never been reported before. The long H3K27me3 regions clustered in four areas, which we defined as H3K27me3 islands, were identified in sorghum. H3K36me3 plays some role in inhibiting the deposition of both H3K27me3 and H2A.Z, which may serve as partial motivation for the removal of H3K27me3 and H2A.Z in leaf and root. All the 7 histone marks are involved in the regulation of tissue-specific genes, especially the specific expression of C4 genes in leaf and peroxidase (POD) encoding genes in root, which are involved in the photosynthesis in leaf and lignin synthesis in root, respectively. These marks except H3K36me3 and H3K27me3 also engage in the regulation of stress genes in response to PEG treatment. However, we found that differential enrichment of histone marks on many tissue-specific genes was observed only between leaf and root but hardly in response to PEG treatment, although expression of these genes changed after PEG treatment.

## Introduction

Histone marks play important roles in gene regulation, and they exhibit different distribution patterns in intragenic regions. Histone acetylation and H3K4me3 tend to mark transcription start site (TSS) regions (a short region immediately downstream from TSS), while H3K36me3 are more likely to be enriched in gene body regions (a longer region from TSS to transcription end site (TES)) (Barski et al. 2007; Guenther et al. 2007; Hu et al. 2020), suggesting that they are involved in governing different processes of gene transcription. Histone acetylation and H3K4me3 have been reported to be involved in transcription initiation by affecting the assembly of pre-initiation complex (PIC), or stimulating the transition from initiation to elongation (Ding et al. 2012; Lauberth et al. 2013; Gates et al. 2017; Wang et al. 2023). H3K36me3, generally considered as a hallmark of transcription elongation, is required for co-transcriptional splicing (Guenther et al. 2007; Iwamori et al. 2016; Sorenson et al. 2016). The distribution patterns of some histone marks are similar in different species while the others are not. For example, in humans and yeast, H3K36me3 is excluded from TSS of genes while there is a clear enrichment peak of the mark around TSS in plants (Barski et al. 2007; Li et al. 2015; Liu et al. 2016; Suzuki et al. 2016). In animals, H3K27me3 can spread to hundreds of kilobases and cover several genes per region (Schwartz et al. 2006). However, most of H3K27me3 regions in *Arabidopsis* are limited to single genes (Zhang et al. 2007). The different distribution patterns of these histone marks may reflect diverse deposition mechanism of them in different species.

The complex crosstalk between different histone modifications is established, since the activity of histone modifiers depositing one histone mark could be either promoted or inhibited by other histone marks. For instance, H2B ubiquitylation is required for simulating the activity of the MLL complex to catalyze H3K4me3 (Hsu et al. 2019). H3K4me3 provides the binding platform for the HAT complexes, facilitating histone acetylations (Taverna et al. 2006; Ringel et al. 2015). The interplay between repressive marks such as H3K27me3 and H2AK119ub1, which are deposited by Polycomb repressive complex 2 (PRC2) and Polycomb repressive complex 1 (PRC1) respectively, have also been unraveled (Blackledge and Klose 2021). Interestingly, both PRC2 and PRC1 can read the two marks to establish or reinforce these modifications (Blackledge and Klose 2021). By contrast, H3K4me3 and H3K36me3 inhibit the activity of PRC2 (Schmitges et al. 2011; Finogenova et al. 2020). Furthermore, it has been demonstrated that H3K36me3 and H3K27me3 are rarely present on the same histone tail in various species (Yuan et al. 2011; Yang et al. 2014), suggesting that the inhibitory effect of H3K36me3 on deposition of H3K27me3 is conserved. Indeed, genome-wide analysis of chromatin states indicates that the distributed regions of some histone marks highly co-exist while the others are seldom overlapped, which is well consistent with the crosstalk between them (Ernst et al. 2011; Sequeira-Mendes et al. 2014).

Sorghum is an ideal model species for analyzing genetic features of C4 plants due to its small genome and natural diploid. Besides, it also exhibits high tolerance to various abiotic stresses, making it a good candidate for studying mechanism of plant stress tolerance. In this paper, genome-wide analysis of various histone modification enrichment was conducted in sorghum root and leave before and after PEG treatment to explore the role of epigenetic marks in the regulation of sorghum development and stress response. We found that distribution pattern of these marks in sorghum was similar to that reported in other plant species. However, sorghum-specific epigenomic features were also revealed.

## Results

### Distribution pattern analysis of histone marks reveals four H3K27me3 islands in sorghum

Three replicates for RNA-seq and two biological replicates for ChIP-seq in leaf and root of sorghum were performed with antibodies against various histone modifications (H3K9ac, H3K27ac, H3K4me3, H3K36me3, H3K4me2 and H3K27me3) and H2A.Z. The three replicates of RNA-seq analysis were highly correlated (Fig. S1a). For most of the ChIP-seq samples, more than 20 million reads were obtained, the overall mapping rate was more than 85% (Table S1), the FRiP value and the SPOT value were greater than 0.5 (Fig. S1b). The two biological replicates of ChIP-seq were also highly correlated (Fig. S1c). Genome-wide view of histone mark enrichment demonstrate that the distribution of all the 7 histone marks was highly correlated with gene density, which is relatively higher at both ends of each chromosome and lower in the middle of the chromosome (Fig. S2). These results suggest that all the data are reliable for subsequent analysis.

To attempt to investigate the distribution pattern of histone marks, intragenic peaks (located from 1.5 kb upstream from gene TSS to 1.5 kb downstream from gene TES) were called for creating metaplots and heat maps (Fig. 1a and 1b). The results showed that H3K9ac, H3K27ac, H3K4me3 were predominantly enriched in TSS regions. H3K4me2 and H2A.Z were distributed either TSS regions or gene body regions. H3K36me3 enrichment extended from TSS in gene body but not to TES. The distribution patterns of these histone marks are similar to those in *Arabidopsis* and rice (Zilberman et al. 2008; Roudier et al. 2009; Li et al. 2015; Hu et al. 2020). We also found strong H3K27me3 enrichment in gene body in both root and leaf. Interestingly, low H3K27me3 enrichment in TSS regions was observed in leaf but not in root (Fig. 1b), which has never been reported in plants before. Besides, genome-wide enrichment of H3K4me2, H3K36me3, H3K27me3 and H2A.Z was decreased in root, suggesting their possible regulatory roles in the fine-tuning of developmental genes (Fig. 1a).

**Figure 1.**
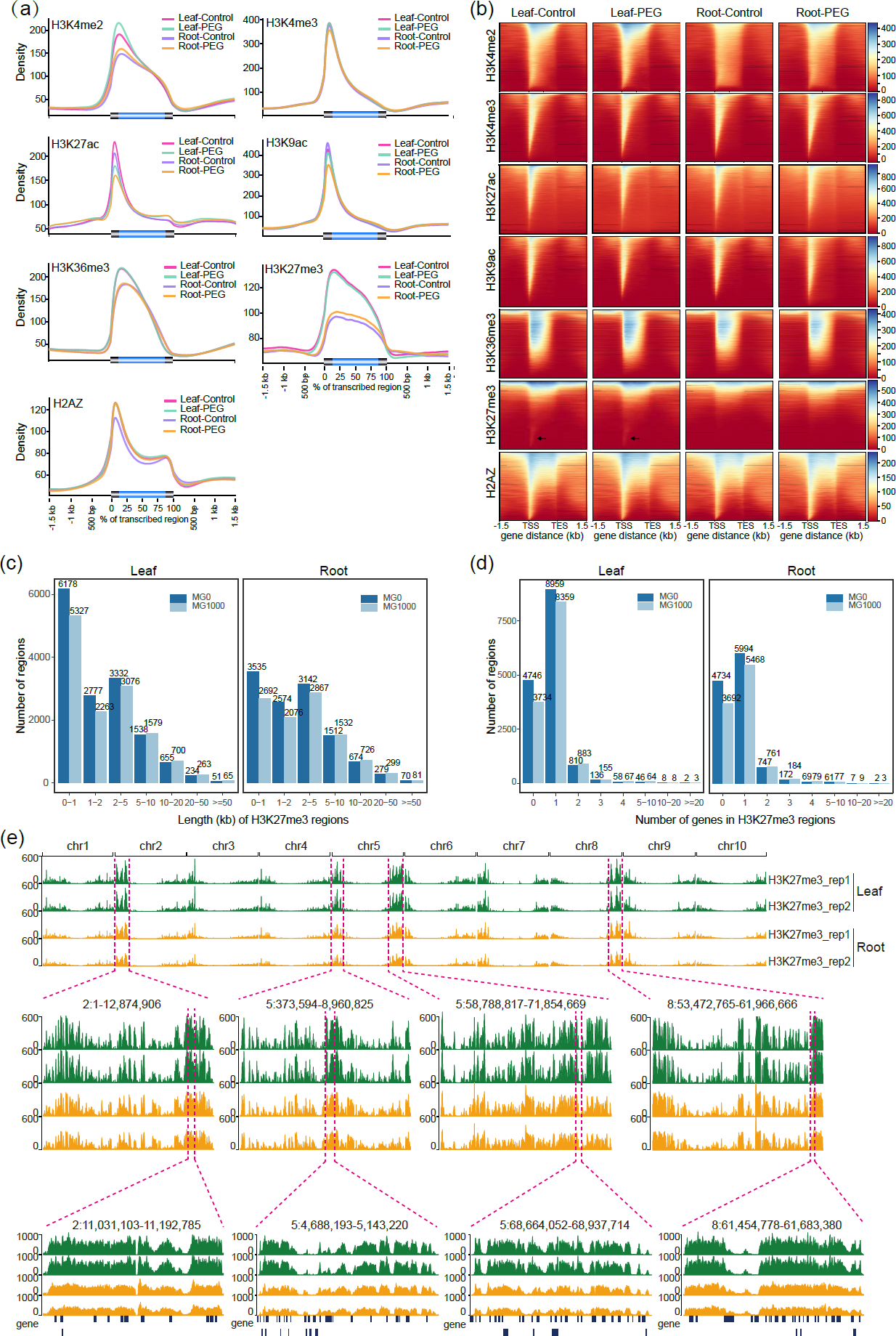
Distribution pattern of histone marks in sorghum. **(a)** Metaplots of 7 histone mark profiles of all genes in leaf and root before and after PEG treatment. **(b)** Heat maps of histone mark enrichment of intragenic region in leaf and root before and after PEG treatment. Black arrows indicate the TSS-enriched H3K27me3 present in leaf but not in root. Two-week-old sorghum seedlings were treated with 20% PEG6000 for 6 hours. Then root and leaf of control (without PEG treatment) and PEG-treated plants were harvested for ChIP-seq. **(c)** Length distribution of H3K27me3 regions in leaf and root. **(d)** The number of genes covered by each H3K27me3 region in leaf and root. “MG1000” indicates that the adjacent H3K27me3 regions were joined when they were separated by less than 1 kb while “MG0” indicates that they were not jointed. **(e)** Genome browser screen shots of 4 H3K27me3 islands on Chromosomes 2, 5, 8. H3K27me3 islands were defined as the areas where a number of long H3K27me3 regions clustered.

In animals, H3K27me3 enrichment can spread to more than one hundred kilobases, which usually cover several genes (Schwartz et al. 2006). It seems that in *Arabidopsis* H3K27me3 coverage is confined to single gene (Zhang et al. 2007). To determine whether H3K27me3 in sorghum is distributed similarly, we analyzed length of H3K27me3 regions and number of genes per region. The results showed that most of H3K27me3 regions were short and covered single genes in sorghum (Fig. 1c and 1d), which is similar as in *Arabidopsis*. However, we found considerable long H3K27me3 regions in sorghum (Fig. 1c and 1d). The maximum length can reach more than two hundred kilobases. Three H3K27me3 regions covered more than 20 genes (Fig. 1d). In addition, there were more short regions and less long regions in leaf than those in root, suggesting that H3K27me3 enrichment in the two organs are reprogrammed. When we visualized genome-wide H3K27me3 regions, we found four areas on Chromosomes 2, 5 and 8 where long H3K27me3 regions clustered, which we termed the H3K27me3 islands (Fig. 1e). The other long H3K27me3 regions scattered in the other part of the genome. The similar analysis was performed in *Arabidopsis* and rice by using the recently deposited high quality data (see Material and methods). The long H3K27me3 regions were also identified in *Arabidopsis* and rice despite that the number was less than that in sorghum (Fig. S3a), especially the regions longer than 50kb. In addition, the H3K27me3 island was not observed in *Arabidopsis* and rice (Fig. S3c). However, the genes covered by long H3K27me3 regions were greater in *Arabidopsis* (Fig. S3b), which may be due to the higher gene density. This indicates that plant epigenomes contain both gene-specific and long region of H3K27me3 enrichment.

### Antagonistic relationship between H3K36me3 enrichment and H3K27me3 enrichment as well as H2A.Z deposition

To analyze the relationship between different histone marks in sorghum, we clustered distribution patterns of each histone mark in intragenic region (from 0.5kb upstream TSS to 3kb downstream TSS) into three categories (C1, C2 and C3), and observe how the other histone marks are distributed in the same category. We found that H3K9ac, H3K27ac, H3K4me3 enriched in the TSS region and H3K36me3 enriched in the gene body region are highly overlapped and associated with actively transcribed genes (Fig. 2a-2d, S4a-S4d). The gene-body-enriched H3K27me3 is exclusive with the above histone modifications especially H3K36me3, and is associated with silent genes (Fig. 2g, Fig. S4g). The partial overlapping of H3K4me3 and H3K27me3 suggests the existence of the bivalent mark in sorghum. The distributions of H3K4me2 and H2A.Z in different regions displayed distinct relationships with the other marks (Fig. 2e-2f, Fig. S4e-S4f). The strong enrichment of H2A.Z in the gene body region (C1) overlapped with the repressive mark (H3K27me3), whereas the weak enrichment of H2A.Z in the TSS region (C3) overlapped with the active marks (H3K9ac, H3K27ac, H3K36me3) (Fig. 2e, Fig. S4e). By contrast, H3K4me2 deposited in the TSS region (C1) co-existed with the repressive mark, whereas that deposited in the gene body region (C2) co-existed with the active marks (Fig. 2f, Fig. S4f). Furthermore, we found that H3K4me2 and H3K4me3 co-existed in many genes although distribution patterns of the two marks in these genes were slightly different. This suggests that the nucleosome simultaneously carrying H3K4me2 and H3K4me3 is common in sorghum, but the deposition of the two marks may be independent.

**Figure 2.**
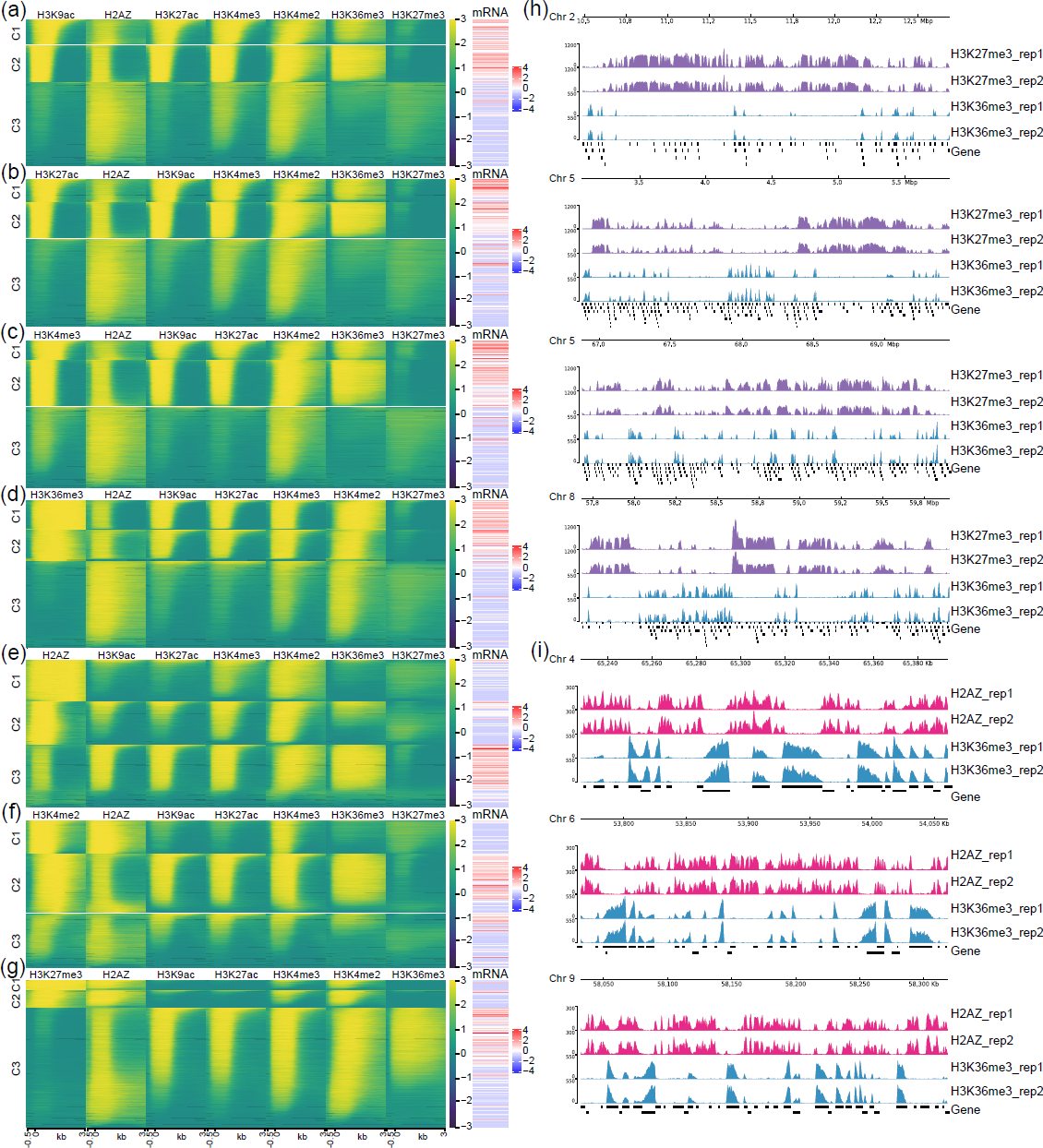
Analysis of overlapping enrichment of different histone marks in sorghum leaf. The enrichment of H3K9ac **(a)**, H3K27ac **(b)**, H3K4me3 **(c)**, H3K36me3 **(d)**, H2A.Z **(e)**, H3K4me2 **(f)**, and H3K27me3 **(g)** in intragenic region were clustered into 3 categories (C1, C2 and C3) respectively and were put on the far left. The other 6 histone mark enrichment and gene expression level were shown next to the clustered marks. **(h)** Genome browser screen shots showing part of the areas in 4 H3K27me3 islands where H3K36me3 and H3K27me3 excluded from each other. **(i)** Genome browser screen shots showing some areas where H3K36me3 and H2A.Z excluded from each other.

Considering the strictly antagonistic distribution of H3K27me3 and H3K36me3, we wonder how H3K36me3 is distributed in H3K27me3 islands. We selected one area in each H3K27me3 island to visualize the distribution of H3K27me3 and H3K36me3. The diagram showed that H3K27me3 and H3K36me3 seldom co-existed at the same gene (Fig. 2h, Fig. S4h), suggesting that they may exclude each other. Indeed, the long H3K27me3 regions are disrupted due to the deposition of H3K36me3. Thus, we speculate that there should be much more and longer H3K27me3 regions than expected in sorghum. As H2A.Z deposition can also extend to generate long H2A.Z regions (data not shown) and H2A.Z is absent in the gene body regions marked by H3K36me3 (Fig. 2e), we randomly selected three long H2A.Z regions and we found that enrichment of H3K36me3 excluded H2A.Z in these regions (Fig. 2i, Fig. S4i). However, the antagonistic relationship between H3K36me3 and H2A.Z is not as rigorous as H3K36me3 and H3K27me3, as at some loci H3K36me3 and H2A.Z co-existed. Taken together, these data suggest that H3K36me3 inhibits the deposition of both H3K27me3 and H2A.Z. To confirm whether the scenario is conserved in plants, we performed the similar analysis in *Arabidopsis* and rice. The results also revealed the antagonistic distributions between H3K27me3 and H3K36me3 as well as H2A.Z and H3K36me3 (Fig. S5a-d).

### Dynamic change of histone marks during development in sorghum

In order to identify which histone marks are more likely to be related with specific gene expression during developmental processes, correlation analysis of differential enrichment of histone marks and gene expression in leaf and root was performed. The metaplots indicated that the overall levels of H3K9ac, H3K27ac, H3K4me3 and H3K36me3 were increased both at root-up-DEGs (Gene expression is up-regulated in roots compared to leaves) in root and leaf-up-DEGs (Gene expression is up-regulated in leaves compared to roots) in leaf (Fig. 3a). The number of genes whose expression changes were positively correlated with changes in these marks was much greater than the number of genes whose expression was negatively correlated with them (Fig. 3b). These supports the positive role of these marks in the regulation of gene expression. In contrast, H2A.Z, H3K4me2 and H3K37me3 levels were significantly decreased at root-up-DEGs in root but slightly decreased at leaf-up-DEGs in leaf (Fig. 3a), and genes negatively regulated by these marks were more than those positively regulated (Fig. 3b), suggesting negative correlation of these marks with gene expression.

**Figure 3.**
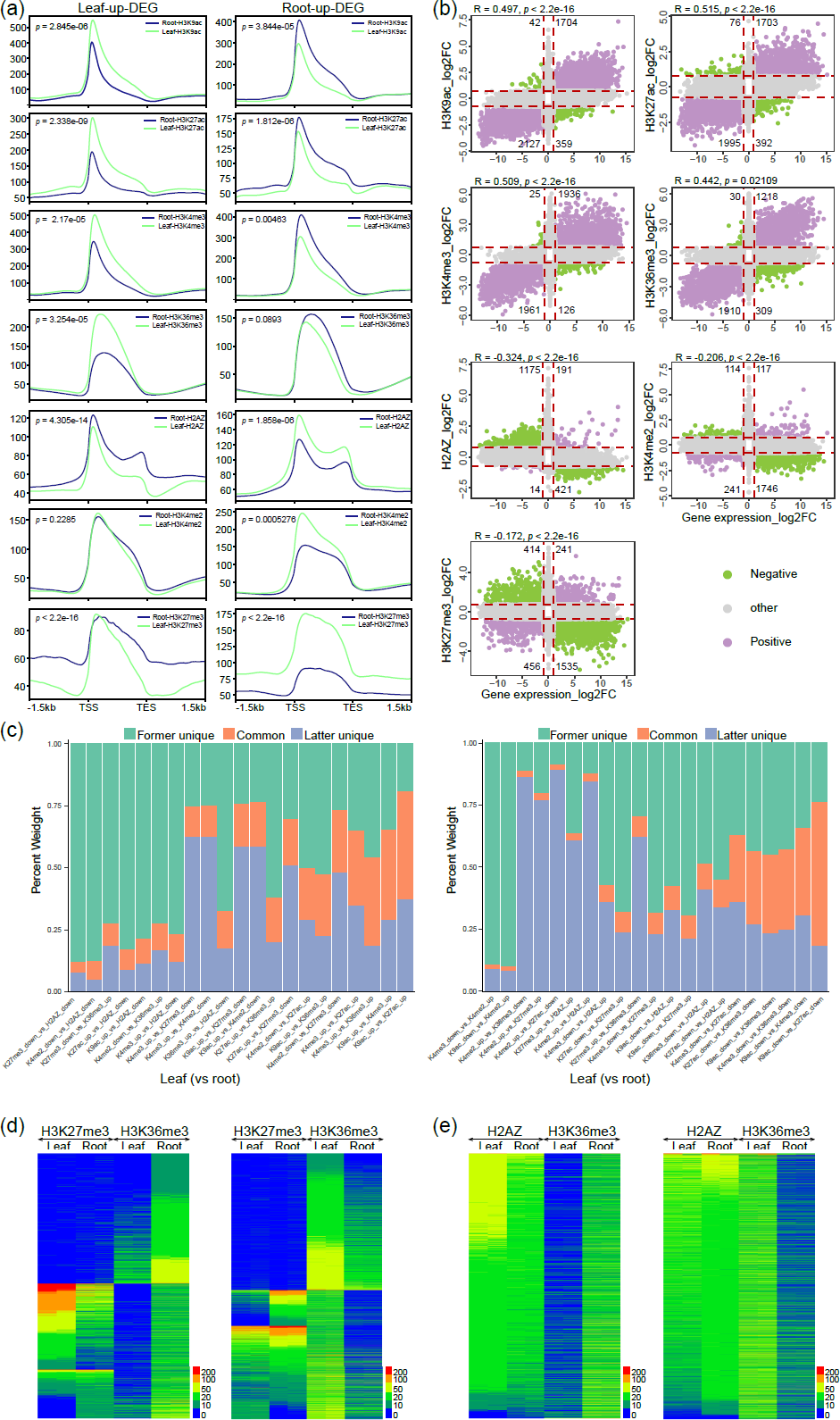
Correlation analysis of differential enrichment of histone marks and differential gene expression in leaf and root. **(a)** Metaplots of 7 histone mark profiles in differentially expressed genes (DEGs) between leaf and root. **(b)** Scatterplots of correlation between change of histone marks (log2FC-root vs leaf) and change of gene expression (log2FC-root vs leaf). Red dots represent the genes with both increased histone mark levels (log2FC > 0.75, p &lt; 0.05) and increased expression levels (log2FC > 1, p &lt; 0.05) and the genes with both decreased histone mark levels (log2FC &lt; −0.75, p &lt; 0.05) and decreased expression levels (log2FC &lt; 1, p &lt; 0.05) (Positive relationship). Blue dots represent the genes with increased histone mark levels and decreased expression levels and the genes with decreased histone mark levels and increased expression levels (Negative relationship). Gray dots represent the genes in which correlation between change of histone marks and change of gene expression was not significant. The number of the co-related genes was indicated. **(c)** The proportion of genes in which two histone marks changes between leaf and root. “Former unique” means the genes in which the former one of the two compared histone mark shown on the horizontal axis change but the latter one does not. “Latter unique” means the genes in which the latter histone mark change but the former one does not. “Common” means the genes in which both histone marks change. **(d)** Heat maps of H3K27me3 and H3K36me3 enrichments in all the genes with changed H3K36me3 (left, H3K36me3 increases in root; right, H3K36me3 increases in leaf). **(e)** Heat maps of H2A.Z and H3K36me3 enrichments in all the genes with changed H3K36me3 (left, H3K36me3 increases in root; right, H3K36me3 increases in leaf).

To investigate how different histone marks cooperate to regulate root-leaf-DEGs (differentially expressed genes between root and leaf), we calculated the proportions of genes that were co-regulated by two histone marks. Considering the opposite roles of different histone marks in gene regulation as described above, the number of genes with increased H3K9ac, H3K27ac, H3K4me3, H3K36me3 or decreased H2A.Z, H3K4me2, H3K27me3 between leaf and root were used for the calculation. The higher proportions of overlapping (more than 30%) were observed between the increase of H3K9ac and the other active marks (H3K27ac, H3K4me3 and H3K36me3) as well as the increase of H3K4me3 and H3K36me3 in both leaf (vs root) and root (vs leaf) (Fig. 3c), suggesting that these histone marks tend to activate gene expression collaboratively. The decrease of H3K4me2 and H3K27me3 also showed a high percentage (30.63%) of overlapping only in leaf (vs root) (Fig. 3c), which demonstrates that they cooperate to repress more gene expression in root. We found that the proportions of overlapping between the increase of H3K36me3 and the decrease of H3K27me3 as well as the decrease of H2A.Z deposition were not as high as expected (8.57% and 15.69% in leaf (vs root), 8.15% and 10.15% root (vs leaf)) (Fig. 3c). This indicates that H3K36me3 enrichment contribute to small part of decline of H3K27me3 enrichment or H2A.Z deposition. In addition, the heat map showed that nearly half of genes with increased H3K36me3 were not marked with H3K27me3 (Fig. 3d). However, most of strong H3K27me3 enrichment were inhibited by the increase of H3K36me3. Similarly, the relatively higher levels of H2A.Z deposition were decreased to moderate levels after H3K36me3 enrichment was increased (Fig. 3e). These results further support the notion that H3K36me3 play part of the roles in the inhibition of H3K27me3 enrichment or H2A.Z deposition.

As shown in Fig. 3a, the differential enrichment of H3K27me3 in root-up-DEGs and leaf-up-DEGs in leaf and root was also obvious outside the gene body region (upstream of TSS and downstream of TES), suggesting that the dynamic change of H3K27me3 could spread outside the intragenic region. Alternatively, H3K27me3 deposition changes in units of long segments instead of single genes. To confirm the suspicion, we examined the number of H3K27m3 peaks with different length differentially enriched in root and leaf and found that although most of differentially enriched H3K27me3 peaks were short (less than 10 kb), more than 300 long H3K27me3 regions (more than 10 kb), consisting of 278 (10-20 kb), 87 (20-40 kb), 12 (40-60 kb), 5 (60-80 kb) and 4 (80-100 kb), were also differentially deposited (Fig. S6a). Genome-wide screenshots indicated that the changes of long H3K27me3 regions scattered in 10 chromosomes and negatively co-related with gene expression in general (Fig. S6b). But close observation on individual regions showed that a number of genes covered by long H3K27me3 regions were not activated despite removal of H3K27me3 (Fig. S6b). For example, two neighboring long H3K27me3 regions decreased in root and localized in chr5:70,880 kb-71,000 kb cover 11 genes, among which only 3 genes were up-regulated (Fig. S6b). This suggests that the alteration of H3K27me3 enrichment in long regions is not always associated with transcriptional activity.

### Leaf-specific expression of photosynthesis genes and root-specific expression lignin biosynthesis genes involves epigenetic regulation

The Kyoto Encyclopedia of Genes and Genomes (KEGG) and Gene Ontology (GO) analysis of root-leaf-DEGs with changed histone marks were performed to discover which pathways were more likely to be subjected to epigenetic regulation. The result showed that leaf-up-DEGs with various changed histone marks were enriched in photosynthesis pathway and localized to chloroplast or plastid. The root-up-DEGs were enriched in Phenylpropanoid biosynthesis pathway and localized to cell wall (Fig. 4a and Fig. S7). For a more detailed analysis of which genes in these pathways were epigenetically regulated, we investigated changes of expression and histone marks of all the photosynthesis genes encoding Photosystem I, Photosystem II, Cytochrome b6/f, ATP synthase, plastocyanin, cytc6 as well as chlorophyll biosynthesis genes. We found that expression of most photosynthesis genes and chlorophyll biosynthesis genes encoding the enzymes that catalyze the reactions from protoporphyrin IX to chlorophyll was dramatically increased in leaf (Fig. 4b). The increased expression of these genes was strongly associated with increased enrichment of H3K9ac, H3K27ac, H3K4me3 and H3K36me3, and partially with decreased enrichment H2A.Z, H3K4me2 and H3K27me3 (Fig. 4b). In particular, dynamic change of H3K27me3 occurred only on a very small proportion of these leaf-specific genes. Besides, a few of these genes were marked with high level of H3K27me3 in root (data not shown), suggesting that H3K27me3 is not required for repression of most of photosynthesis genes in root compared to the other histone marks.

**Figure 4.**
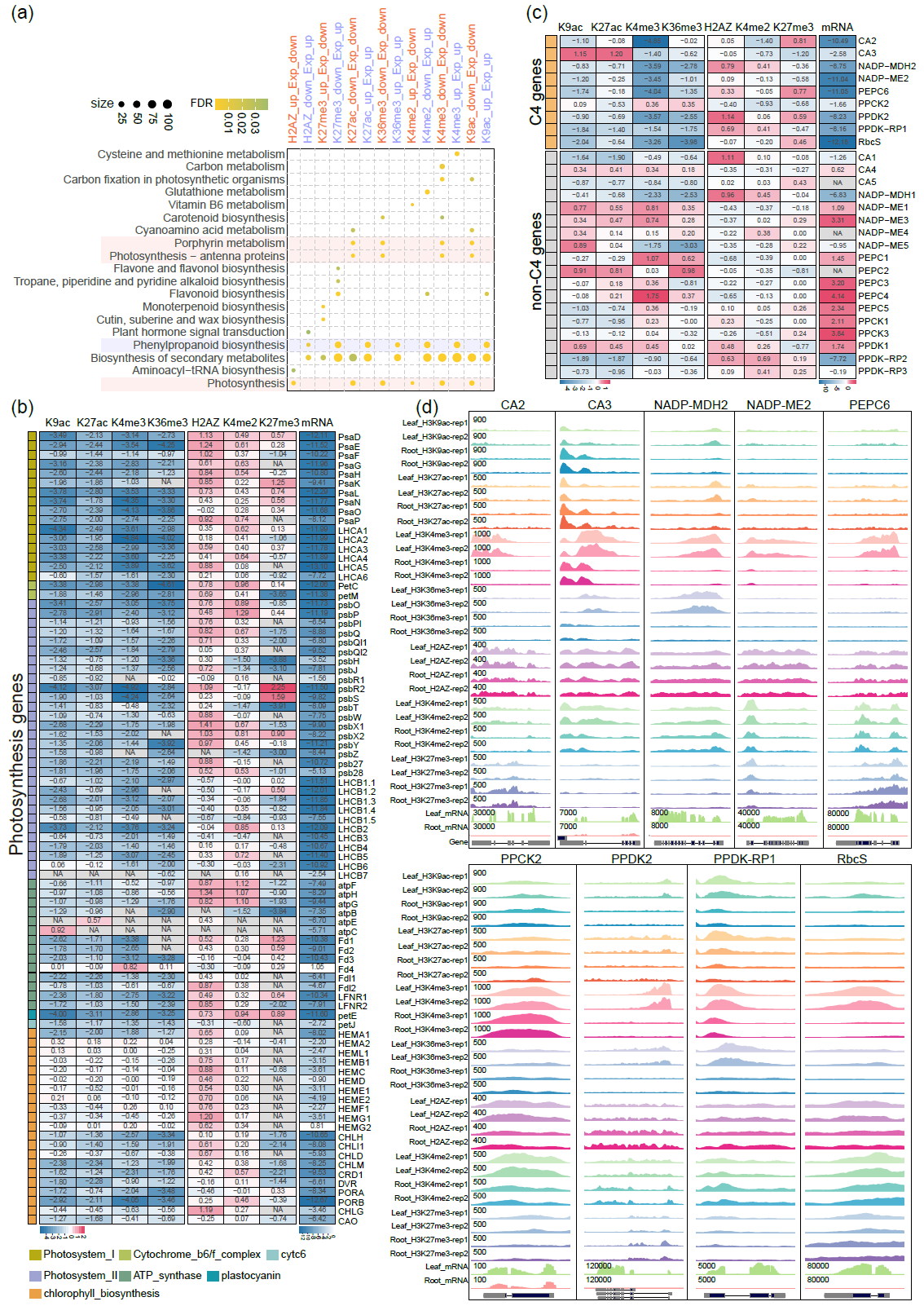
Leaf-specific expression of photosynthesis genes is associated with dynamic change of histone marks. **(a)** KEGG analysis of the leaf-up-DEGs and root-up-DEGs with differential enrichment of various histone marks in leaf and root. The pathways marked with pink background represent leaf-up-DEGs enriched Photosynthesis-related pathways. The pathway marked with purple background represents root-up-DEGs enriched Phenylpropanoid biosynthesis pathway. Heat maps showing fold change (log2FC) of expression and histone marks of photosynthesis genes **(b)** and C4 and non-C4 genes **(c)** in leaf compared to root. **(d)** Genome browser screen shots of histone mark enrichment on C4 genes in leaf and root.

As sorghum is a C4 plant, we would like to learn how C4 genes are regulated by different histone marks. In sorghum there were nine C4 genes: *carbonic anhydrase 2* (*CA2*), *CA3*, *NADP-malate dehydrogenase 2* (*NADP-MDH2*), *nicotinamide adenine dinucleotide phosphate (NADP)-malic enzyme 2* (*NADP-ME2*), *phosphoenolpyruvate carboxylase 6* (*PEPC6*), *PEPC kinase 2 (PPCK2*), *pyruvate orthophosphate dikinase 2* (*PPDK2*), *PPDK regulatory protein 1* (*PPDK-RP1*) and *RbcS* (Tao et al. 2020). We observed that seven C4 genes (except *CA3* and *PPCK2*) exhibited strong leaf expression specificity, and were mostly associated with differential enrichment of H3K9ac, H3K4me3 and H3K36me3 (Fig. 4c and 4d). Among these histone marks, H3K4me3 varied more significantly than the other marks, implying that H3K4me3 acts as the major epigenetic mark for the leaf-specific regulation of C4 gene expression. In addition, similar to the other photosynthesis genes, H3K27me3 was negatively associated with the expression of *CA2* and *PEPC6* only (Fig. 4c and 4d), suggesting a limited regulatory role of H3K27me3 on C4 genes. By contrast, the non-C4 genes homologous to C4 genes showed weaker leaf expression specificity and fewer of them were regulated by dynamic change of histone marks (Fig. 4c). These results suggest that the active histone marks may confer the leaf expression specificity of photosynthesis genes in sorghum.

When we looked into the root-up-DEGs with differentially enriched histone marks in the Phenylpropanoid biosynthesis pathway, we found many of these genes encode lignin biosynthesis enzymes, especially peroxidases (PODs) (data not shown). Therefore, we examined changes of expression and histone marks of all the lignin biosynthesis genes (Fig. 5a). It was shown that for 20 of 52 lignin biosynthesis genes, up-regulation of them in root involves dynamic change of at least one histone mark (Fig. 5b). And the epigenetically regulated genes encode all the enzymes for lignin biosynthesis except C3H, echoing the importance of epigenetic regulation in the lignin biosynthesis in root. More interestingly, we found that vast majority of *POD* genes showed root expression specificity that were associated with most of histone marks (Fig. 5c), although it is not clear how many of them are involved in lignin biosynthesis, especially in root. Particularly, all the 7 histone marks were differentially enriched on four *POD* genes (Fig. 5d). However, It seems that H3K36me3 is not involved in the regulation of most *POD* genes, as they were not marked with H3K36me3 even when they were significantly up-regulated in root (Fig. 5c). Unlike photosynthesis genes, we found decreased enrichment of H3K27me3 on considerable *POD* genes (Fig. 5c), suggesting that repression of these *POD* genes concerns H3K27me3.

**Figure 5.**
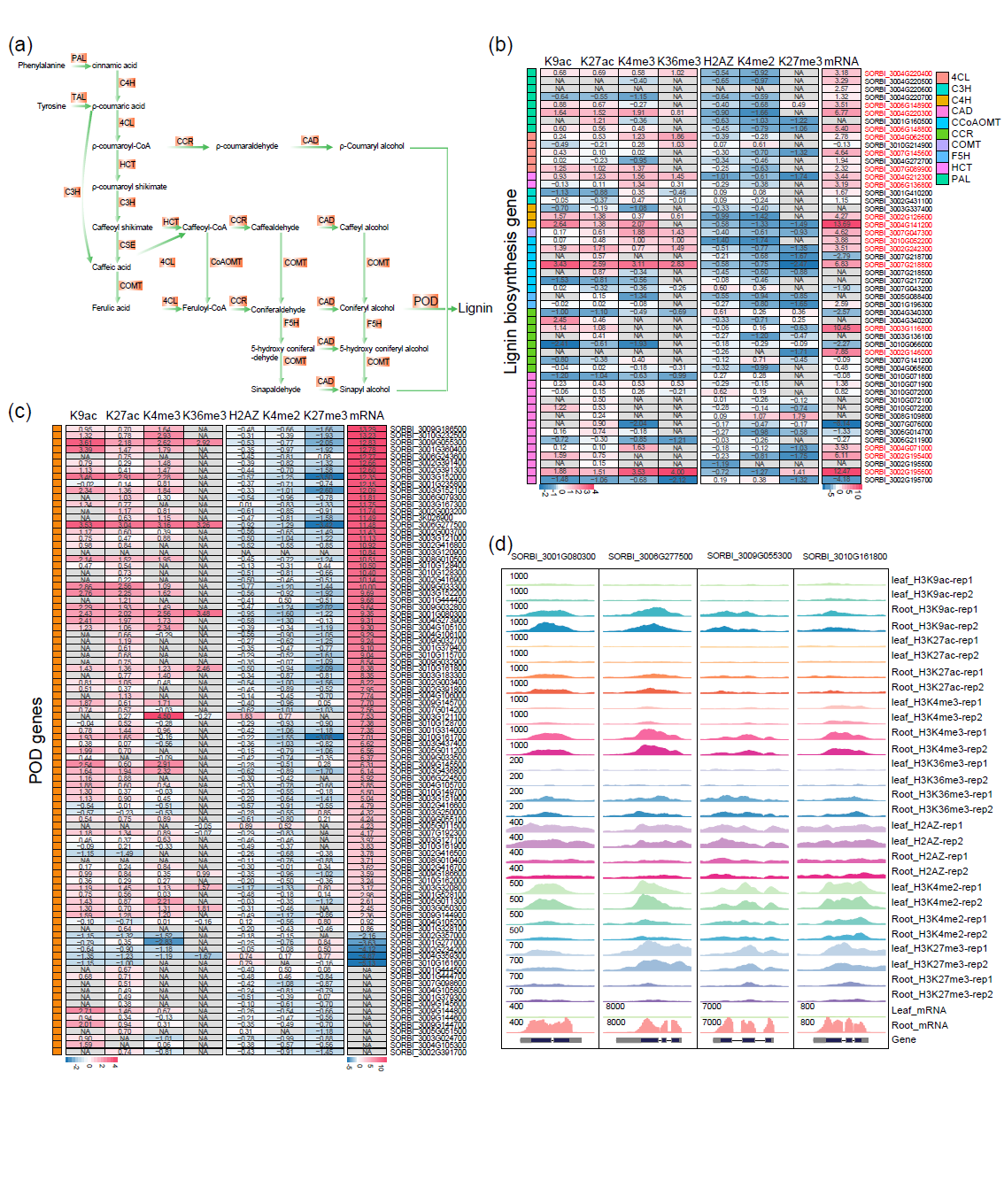
Root-specific expression of lignin biosynthesis and POD genes is associated with dynamic change of histone marks. **(a)** Schematic diagram of lignin synthesis. Heat maps showing fold change (log2FC) of expression and histone marks of photosynthesis genes **(b)** and C4 and non-C4 genes **(c)** in leaf compared to root. **(d)** Genome browser screen shots of histone mark enrichment on ten POD genes with topmost expression change in leaf and root.

### Gene regulation in response to drought stress concerns histone marks except H3K36me3 and H3K27me3

To explore how histone marks engage in the induction of drought-responsive genes, we investigated the levels of histone marks in PEG-up-DEGs and PEG-down-DEGs in response to PEG treatment in root and leaf. The metaplots showed that PEG treatment only slightly affects the overall levels of some histone marks (Fig. 6a and 6b). For example, H3K9ac, H3K27ac, H3K4me3, H3K36me3 levels in PEG-up-DEGs in leaf and H3K36me3 level in PEG-up-DEGs in root showed a little increase. H3K9ac and H3K27ac levels in PEG-down-DEGs in leaf and root was slightly decreased. H2A.Z and H3K4me3 levels in PEG-down-DEGs in leaf and root rose and H2A.Z level in PEG-up-DEGs in leaf and root declined. These results are consistent with the regulatory roles of these marks. Then we performed association analysis of changes of histone marks and gene expression. In both leaf and root, only a few genes had altered H3K36me3 and H3K27me3 responding to PEG treatment (Fig. 6c and 6d), which suggests that these two marks are probably not involved in the PEG-responsive gene expression regulation. H3K4me3 was more likely to be associated with gene activation in response to PEG treatment, as the number of genes with increased H3K4me3 levels and gene expression in leaf (n = 138) and root (n = 363) was far more than the number of genes with the other relationships between H3K4me3 and gene expression (Fig. 6c and 6d). H3K9ac and H3K27ac were associated with either gene activation or repression when the level of these marks was increased or decreased responding to PEG treatment in leaf and root (Fig. 6c and 6d). Surprisingly, for certain number of genes in root but few in leaf changes of H3K9ac and H3K27ac were negatively correlated with changes of gene expression, which seems to contradict to the positive role of these marks. Finally, H2A.Z and H3K4me2 were negatively associated with some gene activation or repression in leaf and root (Fig. 6c and 6d). These results demonstrate that only a small proportion of gene induction in response to PEG treatment require dynamic change of histone marks.

**Figure 6.**
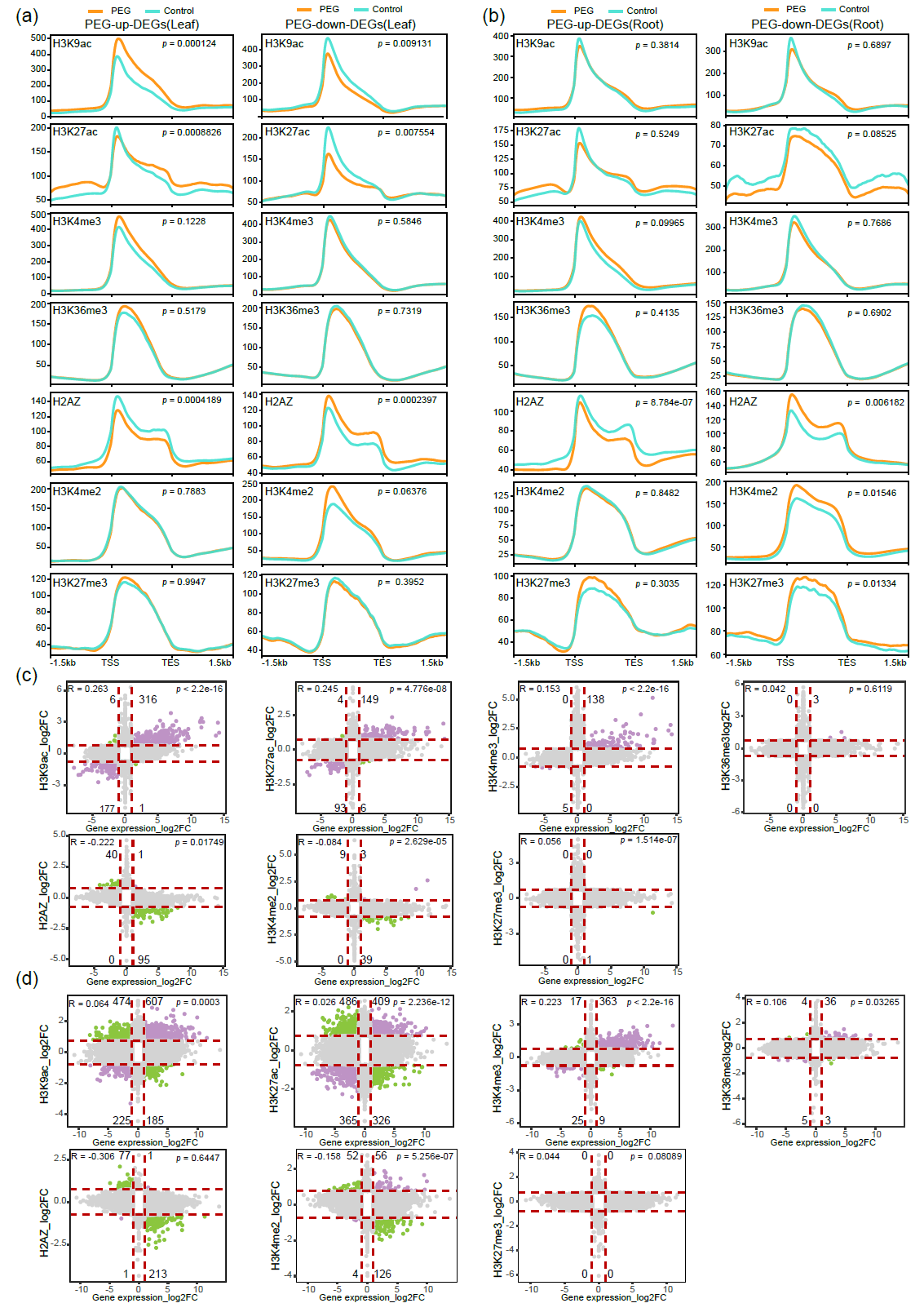
Correlation analysis of changes of histone marks and gene expression in response to PEG treatment in leaf and root. Metaplots of 7 histone mark profiles in differentially expressed genes (DEGs) in response to PEG treatment in leaf **(a)** and root **(b)**. Scatterplots of correlation between change of histone marks (log2FC) and change of gene expression (log2FC) in response to PEG treatment in leaf **(c)** and root **(d)**. Red dots represent the genes with both increased histone mark levels (log2FC > 0.75, p &lt; 0.05) and increased expression levels (log2FC > 1, p &lt; 0.05) and the genes with both decreased histone mark levels (log2FC &lt; −0.75, p &lt; 0.05) and decreased expression levels (log2FC &lt; 1, p &lt; 0.05) (Positive relationship). Blue dots represent the genes with increased histone mark levels and decreased expression levels and the genes with decreased histone mark levels and increased expression levels (Negative relationship). Gray dots represent the genes in which correlation between change of histone marks and change of gene expression was not significant. The number of the co-related genes was indicated.

In order to find out whether histone marks affect the degree of gene induction by PEG treatment, we classified PEG-up-DEGs into three groups: genes without enrichment of histone marks, genes with unchanged histone marks after PEG treatment, genes with changed histone marks after PEG treatment, and investigated fold change of each group gene expression. As PEG-responding gene up-regulation are more associated with increase of H3K9ac, H3K27ac, H3K4me3 and decrease of H2A.Z, the effect of these marks on gene induction was analyzed. We found that the degree of induction of the genes with changed histone marks after PEG treatment was significantly higher than that of the other groups (Fig. S8), suggesting that the dynamic change of histone marks can elevate gene induction to some extent. However, the genes with unchanged H3K9ac, H3K27ac and H3K4me3 were induced to a relatively lower level than the genes without histone marks (Fig. S8). Conversely, the genes with unchanged H2A.Z enrichment were induced to a higher level (Fig. S8). These results demonstrate that the basic level of histone marks is not related to the gene responsiveness.

The direct regulation of many stress genes by transcription factors has been characterized in different plant species such as the genes involved in abscisic acid (ABA) metabolism, signaling and reactive oxygen species (ROS)-related and aquaporins, *late embryogenesis abundant* (LEA), wax biosynthese, polyamine biosynthesis genes (Hu et al. 2022). We found that induction of some of these stress genes also involves epigenetic regulation. For example, *9-cis-epoxycarotenoid dioxygenase 3* (*NCED3)*, the key gene in ABA biosynthesis pathway, and *delta1-PYRROLINE-5-CARBOXYLATE SYNTHASE 1* (*P5CS1)*, the key gene in proline biosynthesis pathway, were up-regulated by PEG treatment in both leaf and root (Table S2). The up-regulation was associated with the increase of H3K9ac, H3K27ac and H3K4me3 (Table S2). Surprisingly, expression of 9 in 10 clade A type 2C protein phosphatases (*PP2C*s) genes, the repressors of ABA signaling, was also increased (Table S2), which usually entails increased enrichment of H3K9ac, H3K27me3, H3K4me3 and decreased enrichment H2A.Z in leaf and root. In addition, epigenetic regulation of many *LEA* genes in response to PEG was also observed (Table S2), although the scenario was different between leaf and root. The expression of these *LEA* genes was induced to a higher degree in leaf where change of epigenetic marks was more significant (Table S2). It is of great interest to ascertain if the higher degree of *LEA* expression is determined by changed epigenetic marks, and whether the differential induction of *LEA* genes in leaf and root has some physiological significance.

### Dynamic change of histone marks is not necessary for the response of tissue-specific genes to drought stress

To learn that for the genes both differentially expressed between leaf and root and responsive to PEG treatment whether the histone marks also change in both conditions (developmentally regulated and PEG responsive), we identified 780 and 2825 Root-up-DEGs that were also PEG-up-DEGs in leaf and PEG-down-DEGs in root respectively, 1359 and 950 Leaf-up-DEGs that were also PEG-down-DEGs in leaf and PEG-up-DEGs in root respectively (Fig. 7a), which were defined as four groups of common DEGs. Then we investigated the number of genes where histone marks changed uniquely in each condition or in both conditions among each of four groups of common DEGs. For the up-regulated genes increase of H3K9ac, H3K27me3, H3K4me3, H3K36me3 and decrease of H2A.Z, H3K4me2, H3K27me3 were analyzed, and for the down-regulated genes decrease of H3K9ac, H3K27me3, H3K4me3, H3K36me3 and increase of H2A.Z, H3K4me2, H3K27me3 were analyzed. The results revealed that for all four groups of common DEGs, the number of genes with change of histone marks uniquely regulated developmentally was much greater than those uniquely responsive to PEG and those in both conditions (Fig. 7b). Moreover, we found some common DEGs where 7 histone marks changed developmentally but none of them changed in response to PEG (Fig. S9). In particular, as we revealed that leaf-specific expression of photosynthesis genes and root-specific expression of lignin biosynthesis genes were associated with dynamic change of histone marks, we attempt to understand how these gene expression was affected by stress. We found that many photosynthesis genes including four C4 genes and lignin biosynthesis genes were slightly down-regulated (Fig. 7c and 7d, Fig. S10), and more than half of *POD* gene expression was decreased to a varying degree after PEG treatment (Fig. 7e). However, histone marks on only a small proportion of these genes were decreased. This indicates that dynamic change of histone marks important for tissue-specific regulation of gene expression is not necessary for the gene induction in response to environmental cues, and some tissue-specific regulatory factors are possibly required for the alteration of histone marks.

**Figure 7.**
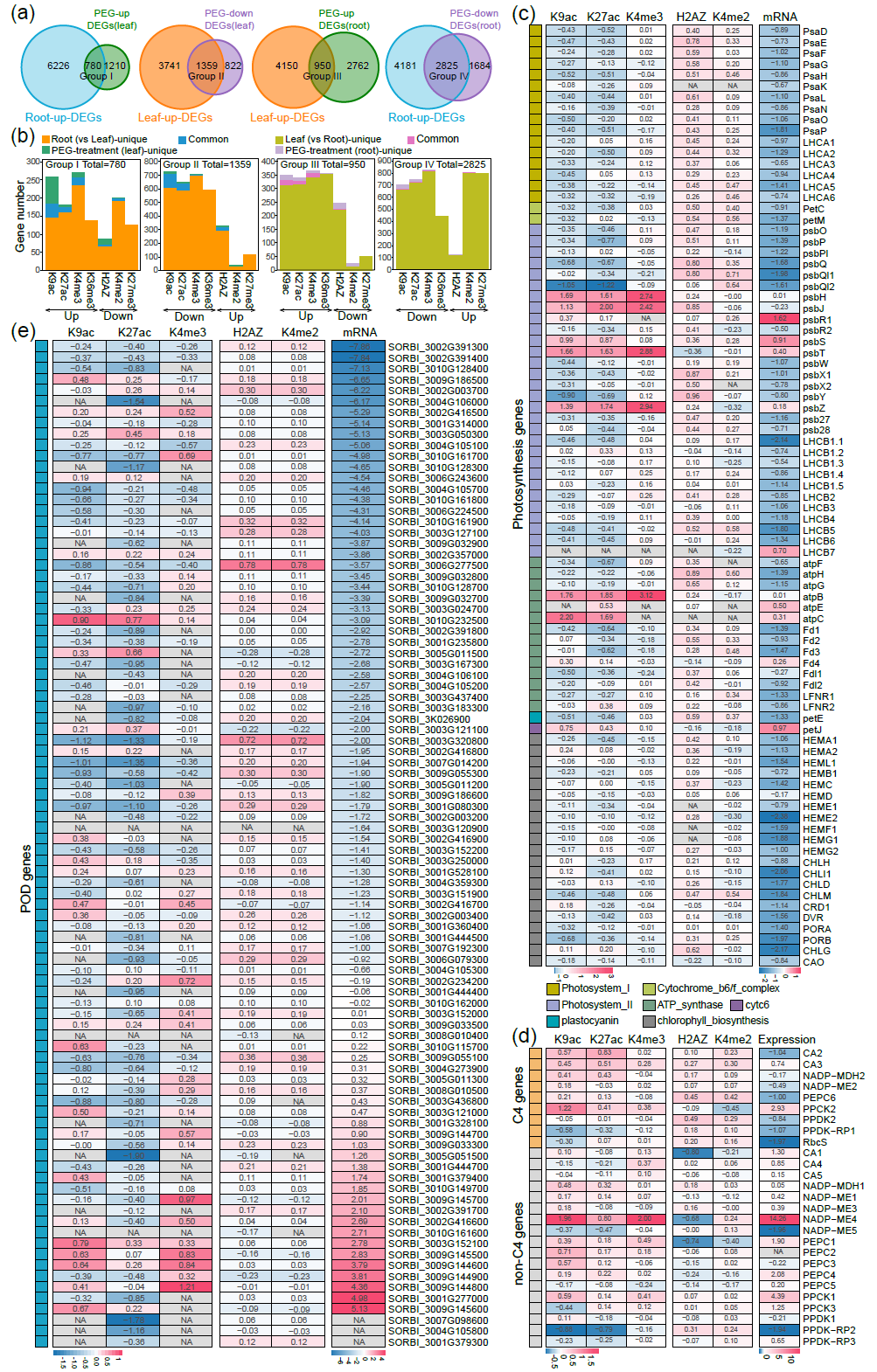
Dynamic change of histone marks in the genes that were both differentially expressed between leaf and root and in response to PEG treatment. **(a)** Venn diagram of overlapping DEGs both between leaf and root and in response to PEG treatment. Four groups (I, II, III, IV) of the overlapping DEGs, also designated as the common DEGs, were identified. Root-up-DEGs: genes with increased expression in root compared to leaf; Leaf-up-DEGs: genes with increased expression in leaf compared to root; PEG-up-DEGs(leaf): genes with increased expression in response to PEG in leaf; PEG-down-DEGs(leaf): genes with decreased expression in response to PEG in leaf; PEG-up-DEGs(root): genes with increased expression in response to PEG in root; PEG-down-DEGs(root): genes with decreased expression in response to PEG in root. **(b)** The overlapping analysis of histone mark changes in four groups of the common DEGs identified in **(a)** both between leaf and root and in response to PEG treatment. “Unique” represents histone marks changed only in this condition but not the other while “common” means histone marks changed in both conditions. The number of “unique” or “common” genes was marked on the column. The total number of each group of the common DEGs was indicated. Heat maps showing fold change (log2FC) of expression and histone marks of photosynthesis genes **(c)** and C4 and non-C4 genes **(d)** before and after PEG treatment in leaf, and POD genes **(e)** before and after PEG treatment in root.

### *Cis*-elements enriched in the promoter of genes with changed histone marks

Through above observations that the responses of chromatin dynamics to developmental or environmental cues were distinct in leaf and root, we considered that tissue-specific factors may direct the change of chromatin structure. Transcription factors (TFs) are the best candidates for the directing factors. *Cis*-elements analysis of promoter of genes (1.5kb upstream from TSS) with changed histone marks was performed to predict which kinds of TFs are responsible for directing the writing or erasing of histone marks in different tissues. The five most enriched *cis*-elements for each variation of histone marks were shown in Fig. 8. For developmental genes, the *cis*-elements of Dof, AT-hook, MADS families TFs were associated with most of histone mark variations (Fig. 8a), suggesting that they were generally involved in the deposition or removal of histone marks in different tissues. The C2H2 Zinc finger TFs binding *cis*-element was inclined to be enriched in the genes with down-regulated histone marks in leaf (Fig. 8a), making it a good candidate as the tissue-specific guider for changing histone marks. In addition, ARID/BRIGHT TFs were responsible for some of histone mark variation (Fig. 8a). More specifically, YABBY TFs were associated with deposition of H3K36me3 in leaf (Fig. 8a). For PEG-induced genes, Dof, AT-hook, C2H2 Zinc finger and AP2 TFs were identified to be the most enriched *cis-*elements binding TFs (Fig. 8b). The bZIP TFs were more likely to be associated with up-regulation of H3K4me3 in leaf (Fig. 8b). MADS box *cis-*element was mostly enriched in the genes with decreased H3K9ac in leaf (Fig. 8b).

**Figure 8.**
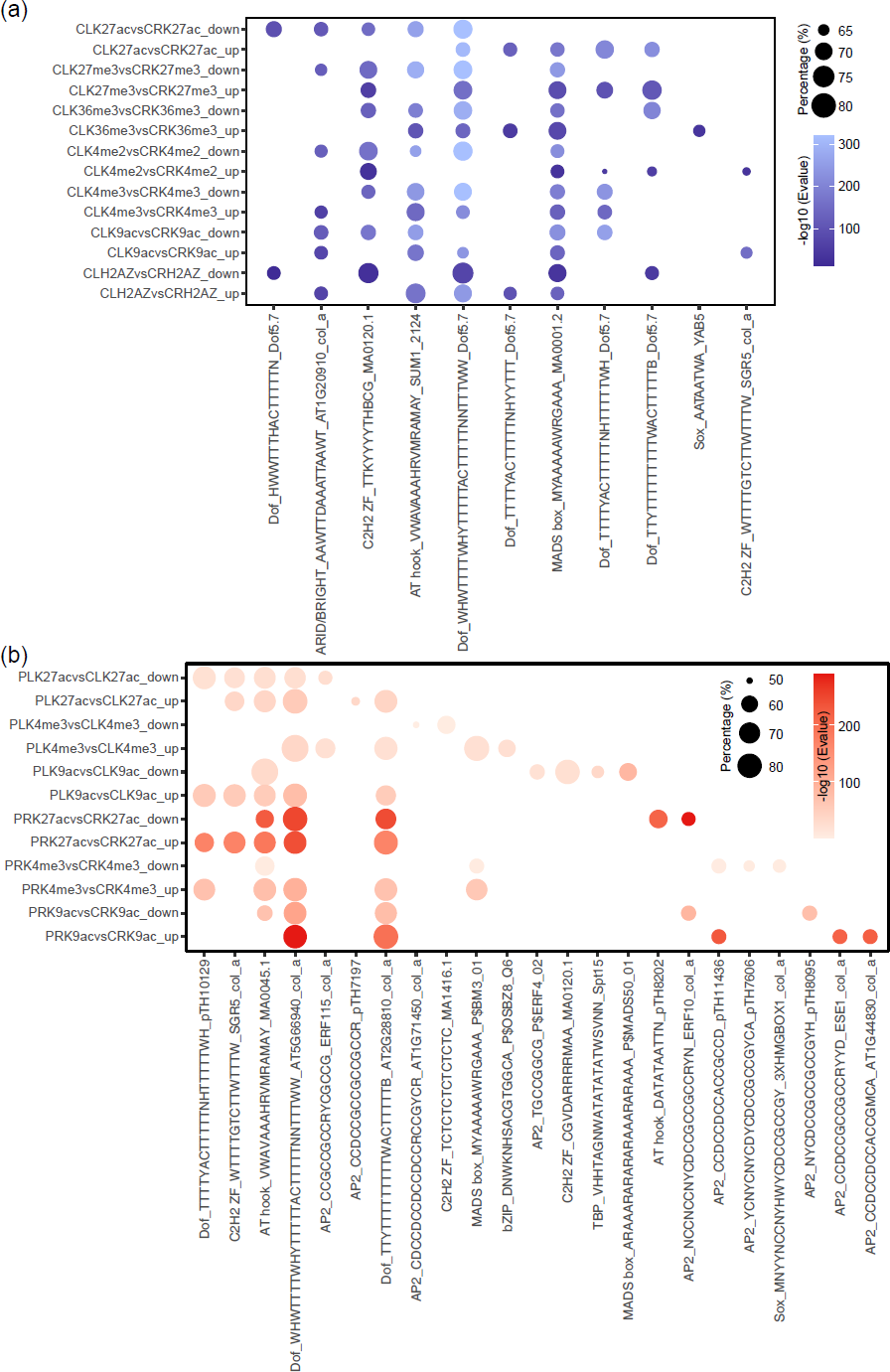
Cis-elements analysis of genes with histone marks changing between leaf and root (a) and in response to PEG treatment (b) CL: Leaf control; PL: Leaf PEG treatment; CR: Root control; PR: Leaf PRG treatment. The percentage (%) of the genes possessing the cis-element relative to the total genes with the histone mark changing was marked on the column.

## Discussion

### The unique feature of H3K27me3 enrichment and function in sorghum

In this study we investigate the genome-wide enrichment of histone marks including H3K9ac, H3K27ac, H3K4me3, H3K36me3, H3K4me2, H3K27me3 and H2A.Z in sorghum. As reported in other plants, the distribution patterns of these marks in intragenic region are conserved in sorghum. For instance, H3K9ac, H3K27ac and H3K4me3 tend to be enriched in the TSS region, and H3K36me3, H3K4me2, H3K27me3 and H2A.Z were distributed in the whole transcribed region (Fig. 1a and 1b). However, early report revealed that H3K27me3 enrichment peaks in *Arabidopsis* was much shorter than that in animals (usually 20-50kb) and limited to single genes mostly (Zhang et al. 2007). But our analysis in *Arabidopsis*, rice and sorghum indicates that there are considerable long H3K27me3 regions in these species despite much less than the short regions (Fig. 2a and Fig. S3a). The maximum length of H3K37me3 region can reach more than 200 kb in sorghum. These results suggest that H3K27me3 enrichment can also spread to a long region covering multiple genes in plants. In Drosophila, spreading of H3K27me3 requires specific *cis*-regulatory sequences, called Polycomb response elements (PREs) and pre-existing H3K27me3 (Margueron et al. 2009). In mammalian, spatially interacted nucleation sites are targeted by PRC2 complex to create H3K27me3-forming Polycomb foci for initiation of H3K27me3 spreading (Oksuz et al. 2018). Indeed, we found four H3K27me3 islands where the long H3K27me3 regions were clustered on three chromosomes of sorghum but not in *Arabidopsis* and rice (Fig. 2c and Fig. S3c). It is intriguing to unveil whether the H3K27me3 islands in sorghum are generated by spreading from PREs as in Drosophila or nucleation sites as in mammalian in the future. It has been disclosed that H3K27me3-mediated chromatin interaction takes place in *Arabidopsis* (Huang et al. 2021; Yang et al. 2022), but whether the interaction of chromatin is the prerequisite for H3K27me3 spreading remains to be elucidated. In our opinion, analysis of 3D genome structure of sorghum to reveal whether the four H3K27me3 islands are interacted is necessary for the next step. Besides, LIKE HETEROCHROMATIN PROTEIN 1 (LHP1), the reader of H3K27me3 in plant, has been reported to be responsible for the spreading of H3K27me3 towards the 3’ end of the gene body in *Arabidopsis* (Veluchamy et al. 2016). It is of great importance to investigate whether the homolog of LHP1 in sorghum is involved in the spreading of H3K27me3 enrichment to a longer region.

### H3K36me3 deposition inhibit both H3K27me3 and H2A.Z enrichment

Inhibition of PRC2 activity by H3K36m3 has been reported in animal and plant by biochemical analysis (Yuan et al. 2011; Yang et al. 2014), but whether H3K36me3 can serve as an active mark to restrict H3K27me3 spreading is not clear. Comparative analysis of genome-wide distribution of the two marks is missing. Here, by comparing genome-wide distribution of 7 histone marks in intragenic regions we found that gene-body-enriched H3K27me3 is depleted in the genes marked with H3K9ac, H3K27ac, H3K36me3 (Fig. 3a, 3b and 3d). But among these active marks only H3K36me3 is distributed in the entire gene body region which is similar to the distribution pattern of H3K27me3 (Fig. 3d and 3g), making it a suitable marker for inhibiting H3K27me3 deposition. When we visualized H3K36me3 enrichment in H3K27me3 island, we obtained mutually exclusive distribution patterns of H3K27me3 and H3K36me3 (Fig. 3h). In the long H3K27me3 regions, a gap would be present where H3K36me3 is deposited, indicating the inhibitory effect of H3K36me3 deposition on H3K37me3 modification. But on the other hand, this also demonstrates that H3K36me3 cannot prevent the spreading of H3K27me3 since H3K27me3 is abundant at both sides of H3K36me3 region. It is of great interest to explore how H3K27me3 spreads over H3K36me3 regions. We also observed that strong enrichments of H3K36me3 were most associated with weak H2A.Z enrichments in the TSS region but hardly associated with gene-body-enriched H2A.Z (Fig. 3d and 3e). This reminds us of the antagonistic relationship between H2A.Z and DNA methylation in *Arabidopsis* (Zilberman et al. 2008). We found that similar to H3K27me3, strong enrichment of H3K36me3 disrupted the long H2A.Z regions (Fig. 3i), documenting that the enrichment of H3K36me3 also inhibit H2A.Z deposition. In mammals, H3K36me2/3 serves as platform to recruit DNMT3A for DNA methylation in the gene body in order for preventing spurious transcription initiation (Yano et al. 2022), but the scenario has not been revealed in plants. It is possible that one of the main functions of H3K36me3 and DNA methylation is to promote gene body H2A.Z eviction in plants. In *Arabidopsis* enrichment of H3K27me3 and deposition of H2A.Z are mutually reinforced (Carter et al. 2018), raising the possibility that the active marks H3K36me3 and DNA methylation can counteract both H2A.Z and H3K27me3. However, our results showed that H3K36me3 variation overlapped with part of changes of H2A.Z and H3K27me3 between leaf and root (Fig. 4c), suggesting that H3K36me3 deposition is not the main cause of removal of H2A.Z and H3K27me3.

### The role of histone marks in the regulation of tissue-specific and stress-responsive genes in sorghum

We compared genome-wide enrichment of histone marks in leaf and root to unravel to what extent do histone marks affect transcriptional reprogramming of tissue-specific genes. The number of differential intragenic enrichments of each histone marks is much less than that of DEGs, suggesting that specific genes were epigenetically regulated in response to developmental cues. A large number of photosynthesis genes including C4 genes and lignin biosynthesis genes were among these genes (Fig. 4b-4d, Fig. 5b). The photosynthesis genes exhibit leaf-specific expression which is consistent with their function. Epigenetic regulation of some C4 genes have been reported in several species including sorghum (Offermann et al. 2006; Heimann et al. 2013; Li et al. 2017). Here we reported differential enrichment of histone marks, particularly H3K4me3, on all C4 genes except PPCK2 in leaf and root of sorghum. But the homologous non-C4 genes were not expressed in a leaf-specific pattern and fewer of them were associated with change in histone marks. This suggest that the introduction of epigenetic marks onto C4 genes during evolution of C4 plants may be critical for their leaf-specific expression pattern. The single cell ChIP analysis is further required to determine whether cell-type-specific expression of C4 genes is conferred by cell-type-specific H3K4me3 enrichment, as C4 genes show mesophyll or bundle sheath cell-type-specificity in leaf. Many lignin biosynthesis genes showed root-specific expression. In sorghum, it has been reported that mutation of *brown midrib 12* (*bmr12*), which encodes caffeic acid O-methyltransferase (COMT) and is one of the key enzymes in lignin biosynthesis, leads to reduced lateral root density and altered root anatomy (Saluja et al. 2021). In this study, we revealed that activation of *bmr12* (SORBI_3007G047300) in root mainly implicated the increase of H3K4me3, H3K36me3 and the decrease of H3K27me3. We also identified many *POD* genes that showed strong root-specific expression pattern and were regulated by various histone marks, although whether they are involved in lignin biosynthesis remains to be elucidated. These results indicate that lignin biosynthesis under epigenetic regulation is important for sorghum root development. However, considering the multiple function of PODs (Cosio and Dunand 2009), it is very likely that they control root development through other pathways instead of lignin biosynthesis. Indeed, the role of POD in root development has been characterized in *Arabidopsis* (Marzol et al. 2022).

The dynamic change of histone marks in response to PEG in leaf and root was also compared in this study. H3K9ac, H3K27ac, H3K4me3, H3K4me2, and H2A.Z are associated with gene induction in varying degree in response to PEG (Fig. 5a-5d). The fold changes of the genes with changed histone marks were greater than those of the genes with unchanged histone marks or without enrichment of histone marks (Fig. S8), suggesting that the variation of histone marks can improve the degree of gene induction. However, H3K36me3 and H3K27me3 enrichments in few DEGs changed both in leaf and root after PEG treatment (Fig. 5c and 5d), suggesting that these two marks are probably not involved in PEG-responsive gene regulation in the process of response in sorghum. The study in moss Physcomitrella patens also showed that H3K27me3 change was not significant in response to drought stress (Widiez et al. 2014). However, in *Arabidopsis* salt priming results in shortening and fractionation of H3K27me3 islands (Sani et al. 2013). The H3K27me3 dynamics responding to salt and cold stress has also been revealed in rice (Zheng et al. 2019; Dasgupta et al. 2022). These demonstrate that different stress signals in different species induce distinct responses of H3K27me3 landscape.

## Materials and methods

### Plant growth conditions and PEG treatment

Sorghum (BTx623 variety) seeds were surface-sterilized and then soaked for germination. The germinated seeds were transferred to the nursery box (1/2 MS) to continue growing in a growth room kept at 28lJ with a 12h light/12 dark cycle. The 14-day-old seedlings were transferred to the nursery box containing 20% PEG6000 that was prepared with 1/2 MS. The seedlings transferred to fresh 1/2MS solution was used as the control. After 6 hours treatment, the third leaves and roots were harvested respectively for subsequent experiments.

### RNA-seq

The leaves and roots were harvested and frozen immediately in liquid nitrogen. Total RNA was extracted using TRNzol Universal Reagent (TIANGEN) following the manufacturer’s instructions. The quality of RNA was measured by gel electrophoresis, Nanodrop analyzer, Labchip and Qubit analyzer respectively. mRNA was purified by using oligo(dT) and then fragmented by incubating in fragmentation buffer. The fragmented mRNA was primed with random hexamer primers and reverse-transcribed with Reverse Transcriptase. After end repair, adenylation, adaptor ligation, purification,PCR amplification, and quality control, sequencing was performed on an Illumina HiSeq system.

### ChIP-seq

Chromatin immunoprecipitation (ChIP) was performed as described in Hu et al. (Hu et al. 2020), In brief, the samples were crosslinked with 1% (v/v) formaldehyde. The nuclei were extracted and the chromatin DNA was broken into 200-500bp fragments by sonication (Soniprep 150, energy 5, 15s on and 1min off for 5 times). The antibodies H3K9ac (07-352, Millipore), H3K27ac (07-360, Millipore), H3K4me2 (07-030, Millipore), H3K4me3 (07-473, Millipore), H3K27me3 (A2363, ABclonal), H3K36me3 (ab9050, abcam), H2A.Z (preparation using antigen “GKGLLAAKTTAAKSAEKDKGKKAPV”) were incubated with Protein A agarose (16-157, Millipore) before mixed with sonicated chromatin. The immunoprecipitated DNA was purified for constructing libraries and sequenced on an Illumina HiSeq system.

### Raw sequencing reads filtering

To obtain high quality clean reads of RNA-seq, ChIP-seq and MNase-seq, the raw sequencing reads were trimmed by Trimmomatic v0.32 (https://github.com/usadellab/Trimmomatic) with parameters of “TruSeq3-PE.fa:2:30:10 LEADING:20 TRAILING:20 SLIDINGWINDOW:4:15 MINLEN:36”.

### Analyses of RNA-seq data

Transcript abundance was quantified using pseudoalignment of high quality clean RNA-seq reads to the reference cDNA sequences and gene models from Sorghum_bicolor_NCBIv3 assembly of the variety BTx623 (McCormick et al. 2018), as implemented in Kallisto (v0.48.0) (Bray et al. 2016). The output pseudoalignment bam files were converted to bigwig format using “bamCoverage” tool of deepTools (v3.5.0) (Ramirez et al. 2016). Differentially expressed genes (DEGs) were identified using DESeq2 v1.22 (Love et al. 2014), genes with TPM value higher than 1 in all the three replicates of each sample were regarded as expressed genes, genes with |log2 (Fold change)| > 1, *P* adjust value < 0.05 and at least expressed in one sample were regarded as DEGs.

### Analyses of ChIP-seq data

High quality clean reads were aligned against the reference Sorghum_bicolor_NCBIv3 genome assembly using Bowtie2 (v2.4.4) (Langmead and Salzberg 2012). Aligned reads with MAPQ < 5 were filtered using SAMtools (v1.9) (Li et al. 2009). Duplicated alignments were removed using Picard (v2.23.9) (http://broadinstitute.github.io/picard/). Before peak calling, the extsize were predicted by phantompeakqualtools (v1.2.2) (Landt et al. 2012). Narrow peaks of H3K9ac, H3K27ac and H3K4me3 ChIP-seq reads densities were identified using MACS2 (v2.2.7.1) (Zhang et al. 2008) with parameters “-f BAM -p 1e-5 --mfold 2 20 --nomodel --extsize $spp_result --to-large”. The input control were used by added parameter “-c”. For broad peaks calling including H3K4me2, H3K27me3, H3K36me3 and KH2A.Z, the “--broad” parameter was added. The peaks from biological replicate samples were merged by BEDTools (v2.27) (Quinlan and Hall 2010), two replicate bam files were merged by SAMTools. We then extracted the reads from the merged bam files that mapped on the merged peaks using BEDtools, and converted the output new bam files to RPKM normalized bigwig files by deepTools. The metaplot and heat map profiles of epigenetic and genomic features were generated and plotted by deepTools.

The signal portion of tags (SPOT) and the fraction of reads in called peak regions (FRiP) value were calculated using Hotspot v4.1.0 (https://github.com/rthurman/hotspot) and commands in-house. To assess the repeatability between biological samples, we calculated the correlation coefficients between samples using deepTools. The visualization of reads coverage is available on http://crispr.hzau.edu.cn/cgi-bin/CRISPR-Cereal/gb2/gbrowse/Sorghum_bicolor/.

### Visualization of overall signals

All types of signals were visualized using Circos (version 0.69) (Krzywinski et al. 2009) with the commands available on github (https://github.com/hcph/Sorghum-omics-dataset-analysis).

### Analyses of H3K27me3 regions

The H3K27me3 peaks in bed files and the signal normalized bigwig files were downloaded from the ChIP-hub databases (Fu et al. 2022) using the sample ID of SRX7734769 and SRX7734770 for *Arabidopsis*, and SRX7426661 and SRX7426662 for rice. Peaks in the two replicates with 1 bp overlap (MG0) as well as overlap regions lower than 1,000 bp (MG1000) were merged using BEDTools. The number of H3K27me3 regions with different length distributedwere visualized. To investigate the distribution of H3K27m3 regions with different number of overlapped genes, we firstly found the overlapped genes for each H3K27me3 region, then the number of overlapped genes was calculated. The *Arabidopsis* and rice H3K27me3 regions longer than 10 kb were extracted and the H3K27me3 signal located in these regions were visualized using pyGenomeTracks v3.5 (Lopez-Delisle et al. 2021).

### Detection of differentially histone modified genes (DMGs)

Reads mapped to genes were extracted as described before. The reads density on each genes was calculated by Subread v2.0.0 (Liao et al. 2013). The TPM value was used to quantified the reads density on each gene, followed by DMGs detection using DeSeq2. Genes with TPM value higher than 10 in the two replicates were regarded as modified genes, and genes with |log2 (Fold change)| > 0.75, *P* adjust value < 0.05 and at least being modified in one sample were regarded as DMGs. The pearson correlation coefficients for the differentially modification and expression levels was calculated. Long (>10 kb) H3K27me3 signals located in differentially H3K27me3 modified regions were extracted with BEDtools as described before, the extracted H3K27me3 signal and the corresponding gene expression signal were visualized with Circlize v0.4.15 (Gu et al. 2014) .

### Analyses of relationship between different histone modifications

The signal matrix of the seven histone modification marks was built by deepTools using the merged and RPKM normalized bigwig files as input. The signal of a single mark was used in turn as a template for clustering the signals of the remaining marks into three groups. The datasets in rice (Zheng et al. 2019; Lu et al. 2020; Zhao et al. 2020) and *Arabidopsis* (Carter et al. 2018; Xi et al. 2020) were downloaded from ChIP-hub and analyzed in the same way as in sorghum. The signals located in long peaks (> 10 kb) were extracted and visualized as described before.The number of DMGs with two differential types of histone modifications or only differentially in one histone modifications were calculated and visualized.

### GO and KEGG enrichment analysis

GO and KEGG enrichment was performed using g:Profiler (https://biit.cs.ut.ee/gprofiler/gost), the enriched GO terms and KEGG pathways were visualized using R package ggplot2 v3.3.5.

### Motif enrichment

The promoter sequences (upstream 1,500 bp) of DMGs were extracted using BEDTools. The sequences were submitted into the CentriMO (https://meme-suite.org/meme/tools/centrimo) online analysis toolkit for motif enrichment analysis. The “anywhere” mode was selected to perform local motif enrichment and the motif database was supported by CIS-BP v2.0 (Weirauch et al. 2014).

## Declarations

### Ethics approval and consent to participate

Not applicable

### Consent for publication

Not applicable

### Availability of data and materials

The ChIP-seq, MNase-seq and RNA-seq data generated in this article were deposited in the NCBI Sequence Read Archive (SRA) (http://trace.ncbi.nlm.nih.gov/Traces/sra/) under the BioProject accession number PRJNA952350.

### Authors’ contributions

The project was conceived by YH and supervised by XS. YH performed ChIP-seq, MNase-seq and RNA-seq with help from QH, GZ, XQ and ZD. CH performed the bioinformatics analyses with help from YS and SB. The manuscript was drafted by YH and XS and revised by all the authors. ZH participate in the critical discussion. The authors read and approved the final manuscript.

## Acknowledgements

We sincerely thank the computing platform of the National Key Laboratory of Crop Genetic Improvement in HZAU for providing the computational resources.

---

Figure S1. The quality of ChIP-seq and RNA-seq data obtained in this study.

Figure S2. The overview of genome-wide enrichment of histone marks in sorghum.

Figure S3. Long H3K27me3 regions in Arabidopsis and rice.

Figure S4. Analysis of overlapping enrichment of different histone marks in sorghum root.

Figure S5. The relationship between H3K36me3, H3K27me3 and H2A.Z in Arabidopsis and rice.

Figure S6. Long H3K27me3 peaks change between leaf and root

Figure S7 Gene Ontology (GO) analysis of the leaf-up-DEGs and root-up-DEGs with differential enrichment of various histone marks in leaf and root.

Figure S8. The role of histone marks in the induction of PEG-responsive genes.

Figure S9. Genome browser screen shots of histone mark enrichment in some common DEGs (identified in Figure 6) with 7 histone marks changing between leaf and root but not in response to PEG treatment.

Figure S10 Heat maps showing fold change (log2FC) of expression and histone marks of lignin biosynthesis genes

Table S1 Total reads and mapping rate of ChIP-seq data

Table S2 Changes of expression and histone marks of stress genes

