## Supplementary material for "Dynamics of the epigenetic landscape during development and in response to drought stress in sorghum": Supplementary figures.pdf

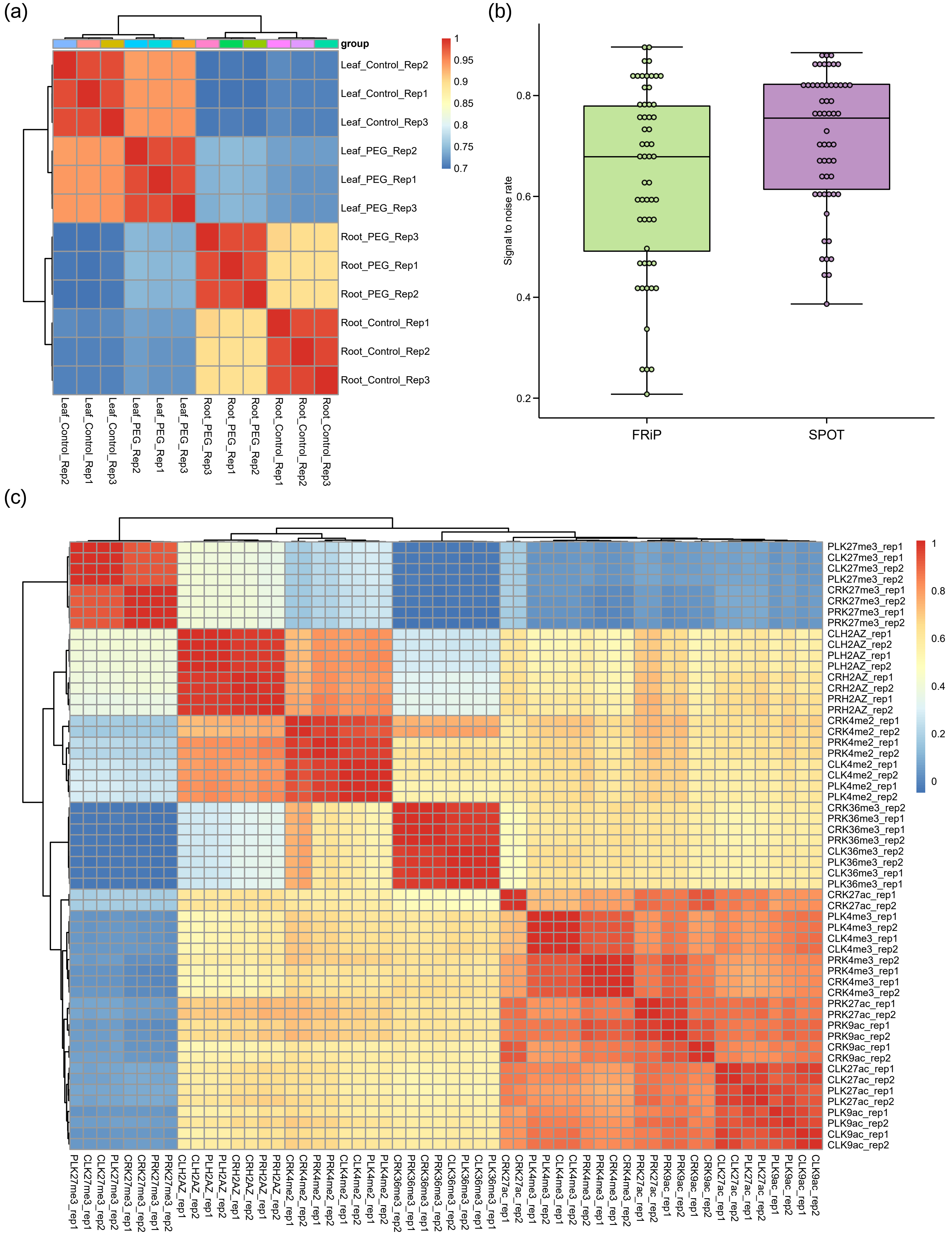


**Figure S1. The quality of ChIP-seq and RNA-seq data obtained in this study.** (**a**) Correlation analysis of three replicates of RNA-seq data. (**b**) The FRiP and STOP values of ChIP-seq data. (**c**) Correlation analysis of two replicates of ChIP-seq data.

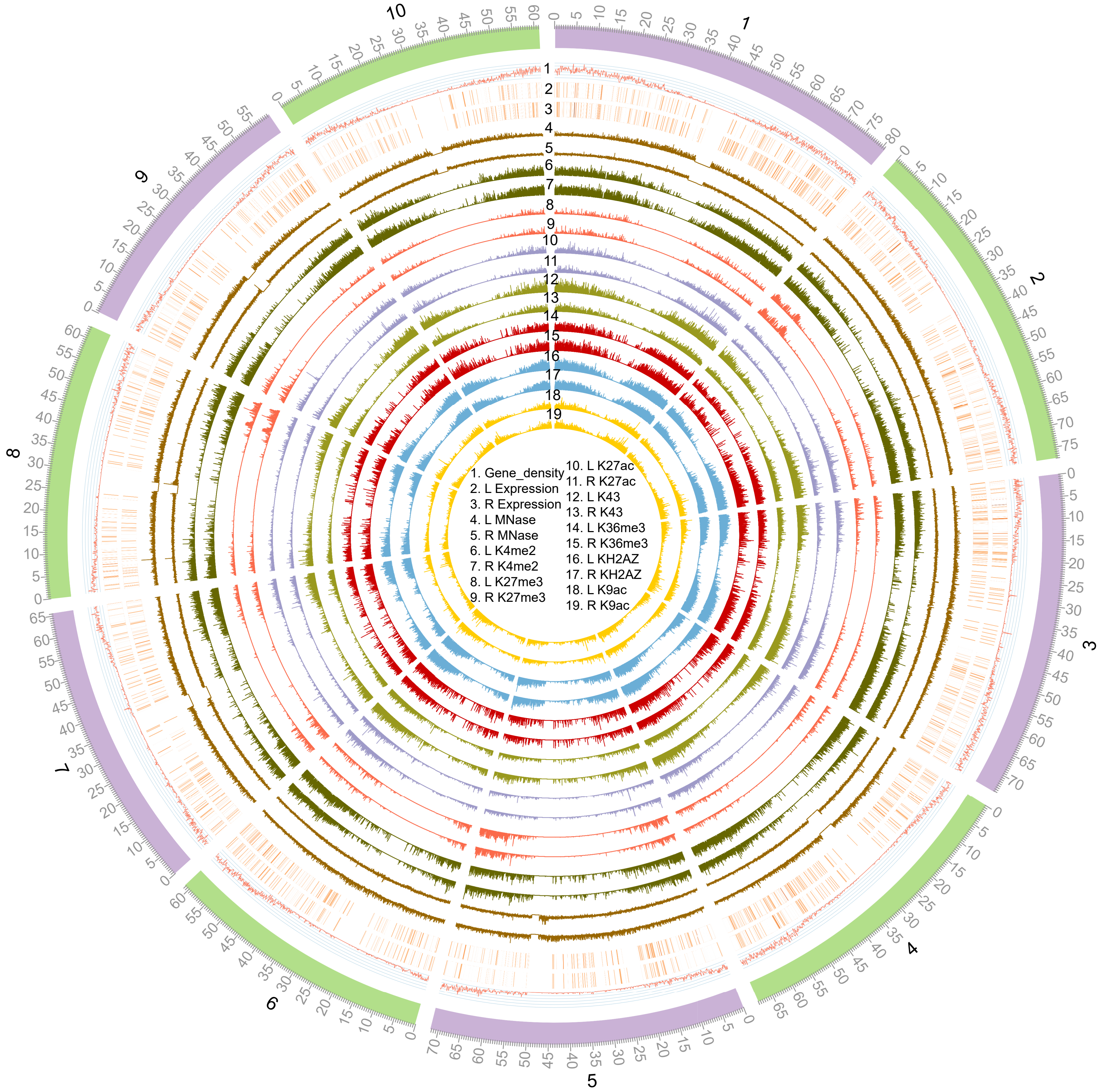


**Figure S2. The overview of genome-wide enrichment of histone marks in sorghum.** L: leaf; R: root.

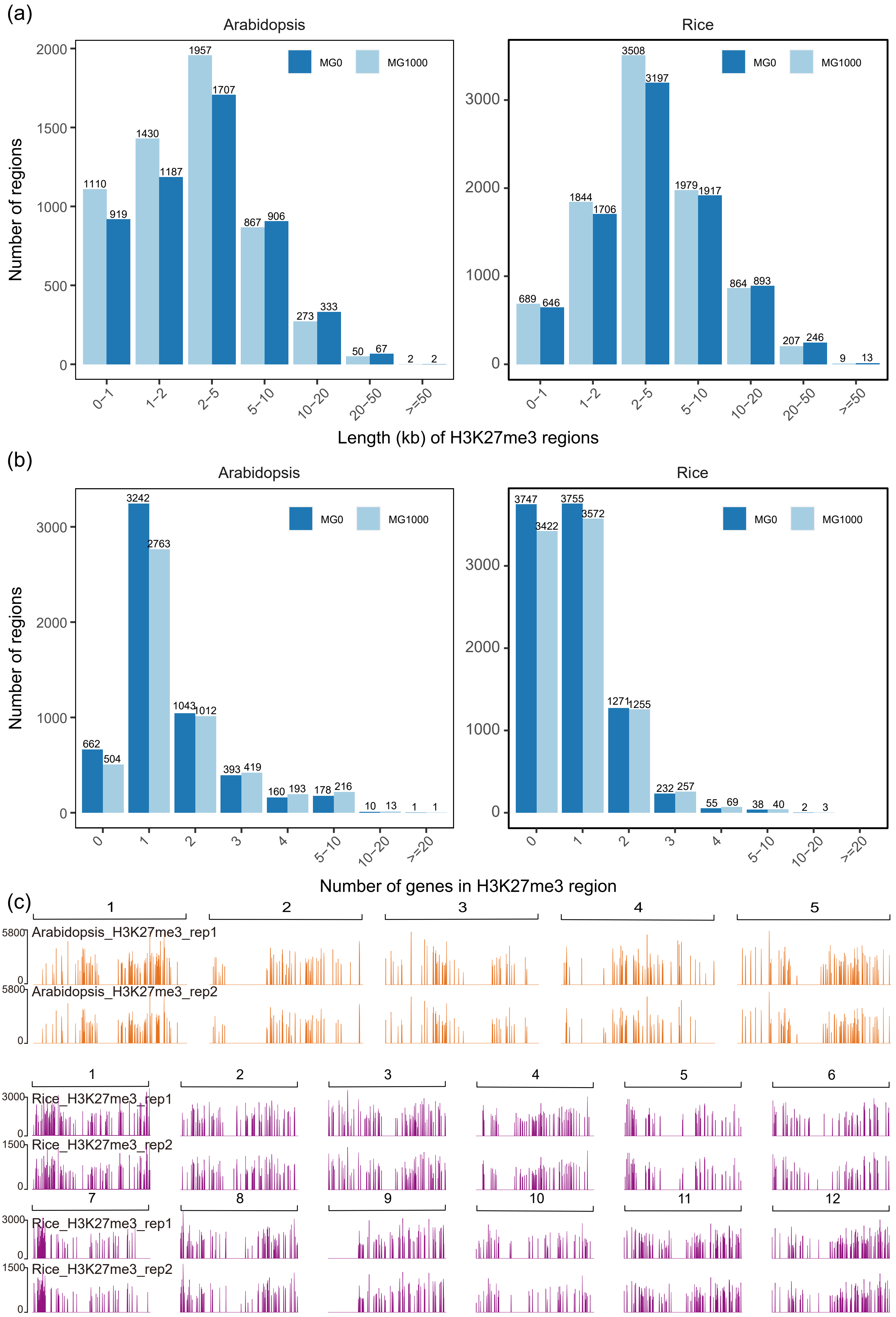


**Figure S3. Long H3K27me3 regions in Arabidopsis and rice.**
(**a**) Length distribution of H3K27me3 regions in Arabidopsis and rice. (**b**) The number of genes covered by each H3K27me3 region in Arabidopsis and rice. “MG1000” indicates that the adjacent H3K27me3 regions were joined when they were separated by less than 1 kb while “MG0” indicates that they were not jointed. (**b**) Genome browser screen shots of genome-wide H3K27me3 enrichment in Arabidopsis and rice.

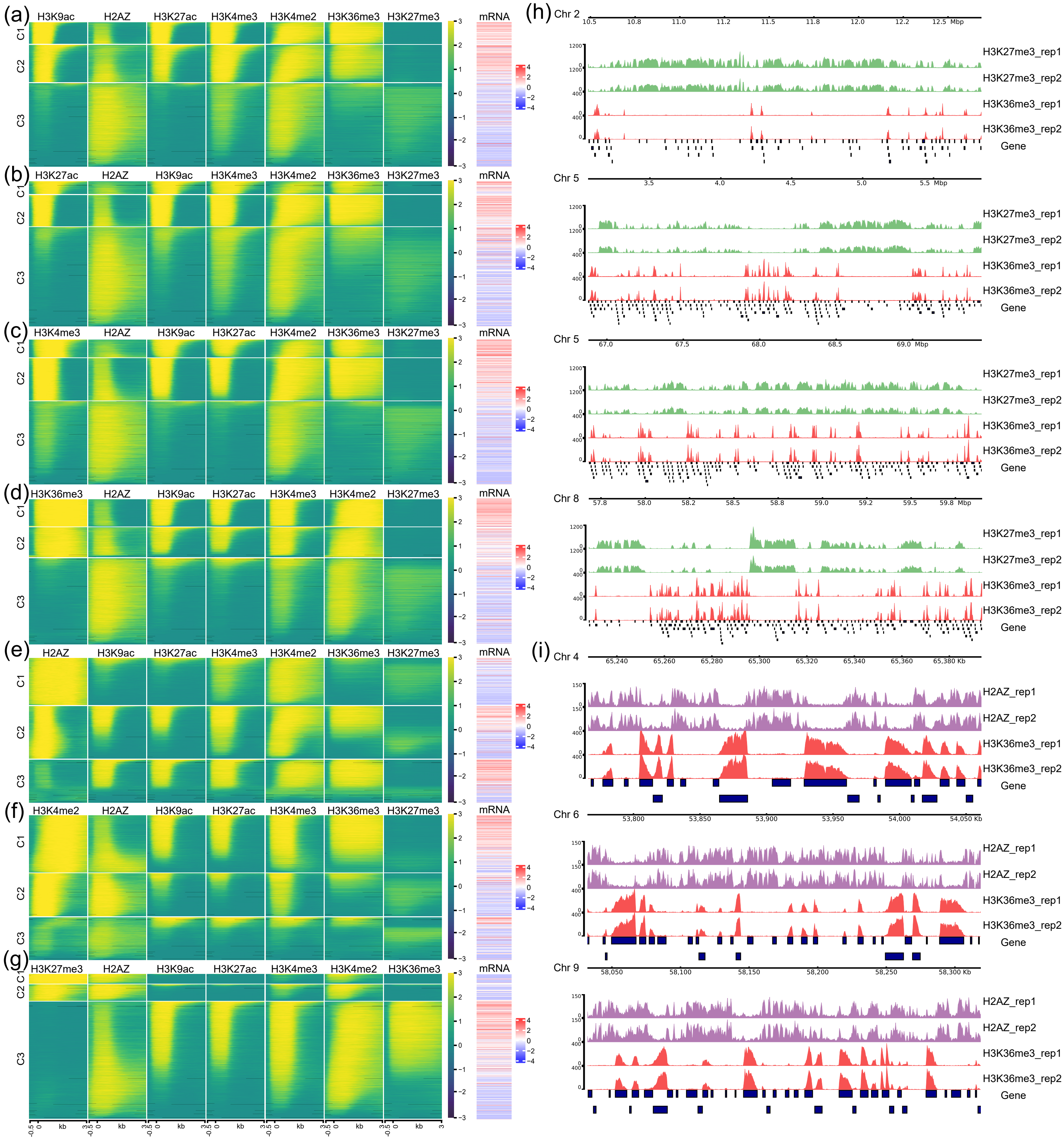


**Figure S4. Analysis of overlapping enrichment of different histone marks in sorghum root.** The enrichment of H3K9ac (**a**), H3K27ac (**b**), H3K4me3 (**b**), H3K36me3 (**d**), H2A.Z (**e**), H3K4me2 (**f**), and H3K27me3(**g**) in intragenic region were clustered into 3 categories (C1, C2 and C3) respectively and were put on the left. The other 6 histone mark enrichment and mRNA level were shown next to the clustered marks. (**h**) Genome browser screen shots showing part of the areas in 4 H3K27me3 islands where H3K36me3 and H3K27me3 excluded from each other. (**i**) Genome browser screen shots showing some areas where H3K36me3 and H2A.Z excluded from each other.

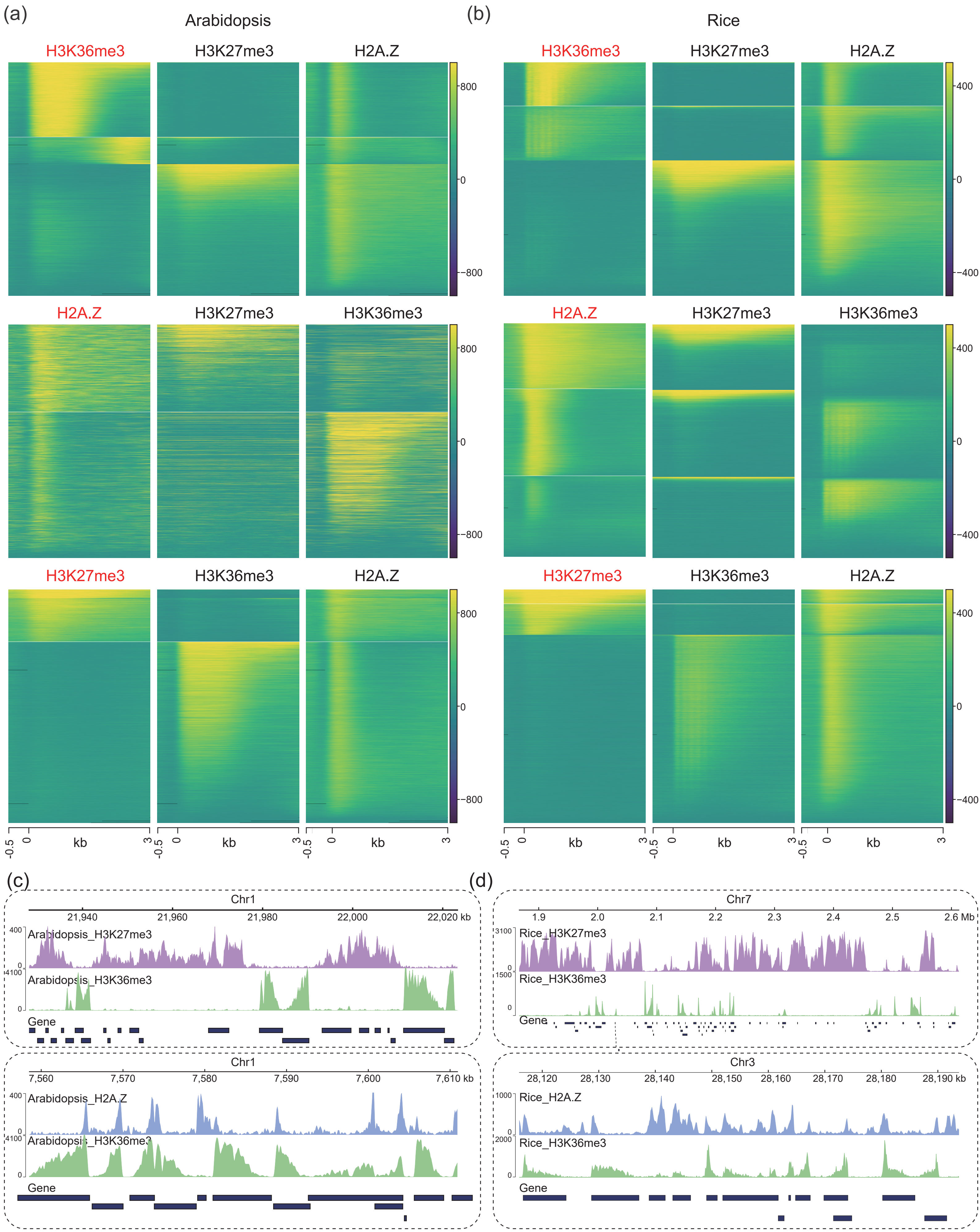


**Figure S5. The relationship between H3K36me3, H3K27me3 and H2A.Z in Arabidopsis and rice.** In Arabidopsis (**a**) and rice (**b**), the enrichment of H3K36me3, H2A.Z and H3K27me3 in intragenic region were clustered into 3 categories respectively and were put on the left. The other 2 histone mark enrichment were shown next to the clustered marks. Genome browser screen shots showing some areas where H3K36me3 and H3K27me3 as well as H3K36me3 and H2A.Z excluded from each other in Arabidopsis (**c**) and rice (**d**).

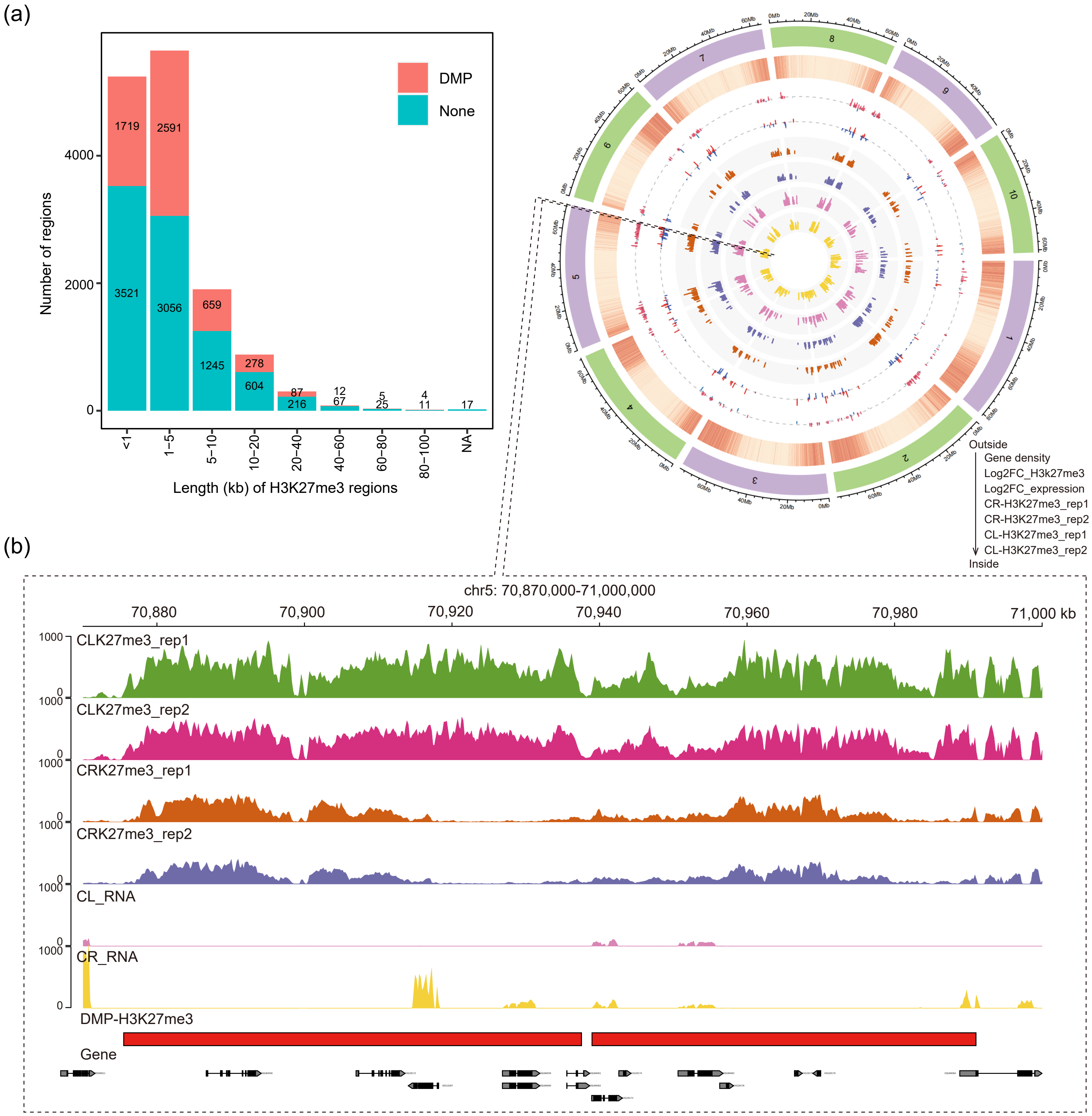


**Figure S6. Long H3K27me3 peaks change between leaf and root** (**a**)Changes of H3K27me3 peaks with different lengths. (**b**)Genome-wide view of change of long H3K27me3 peaks (>10kb) and expression change of genes within these peaks.

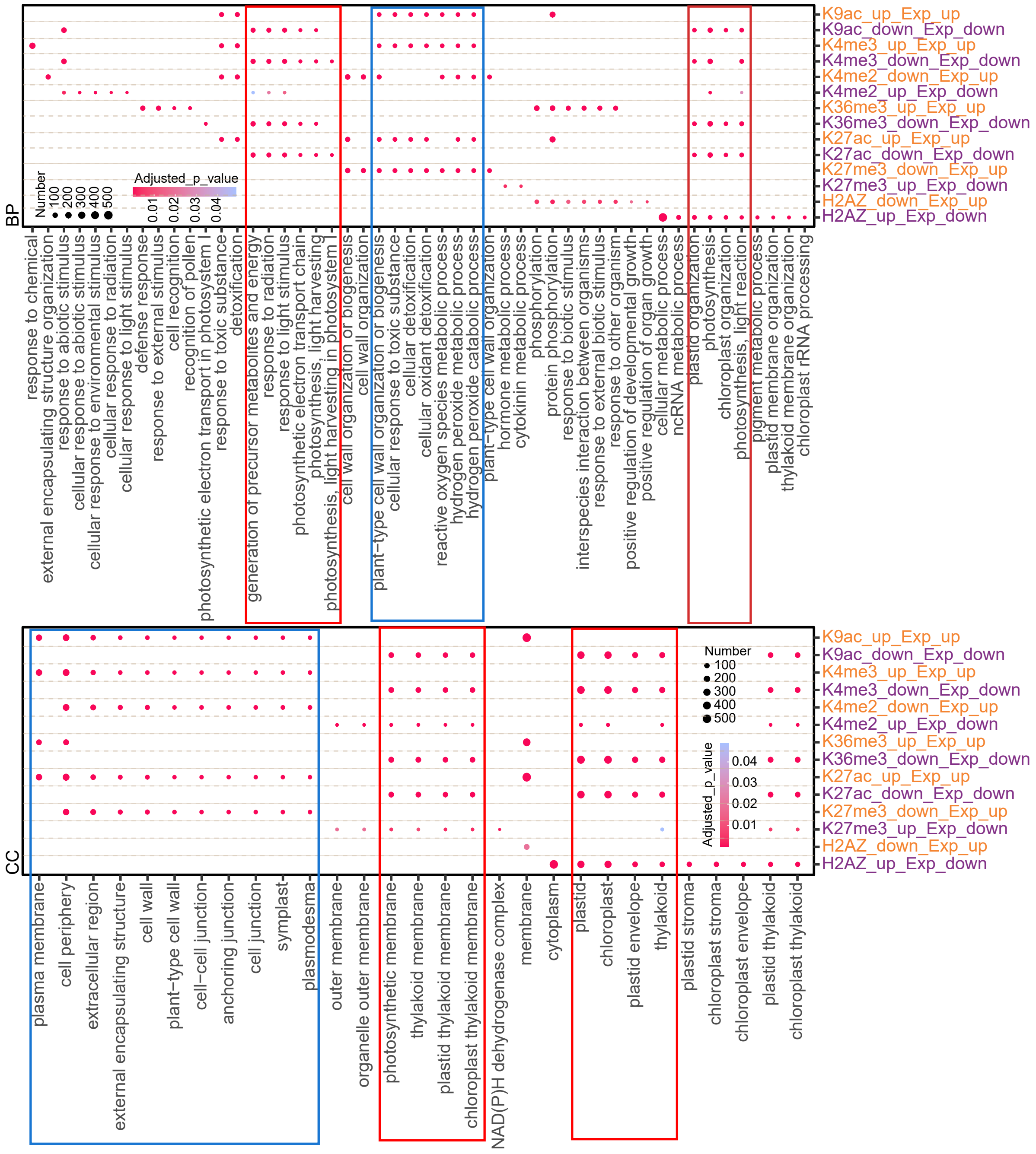


**Figure S7 Gene Ontology (GO) analysis of the leaf-up-DEGs and root-up-DEGs with differential enrichment of various histone marks in leaf and root.** Red rectangle boxes represents leaf-up-DEGs is enriched in Photosynthesis-related pathway. Blue rectangle box represent root-up-DEGs is enriched in cell wall or peroxidase-related pathway.

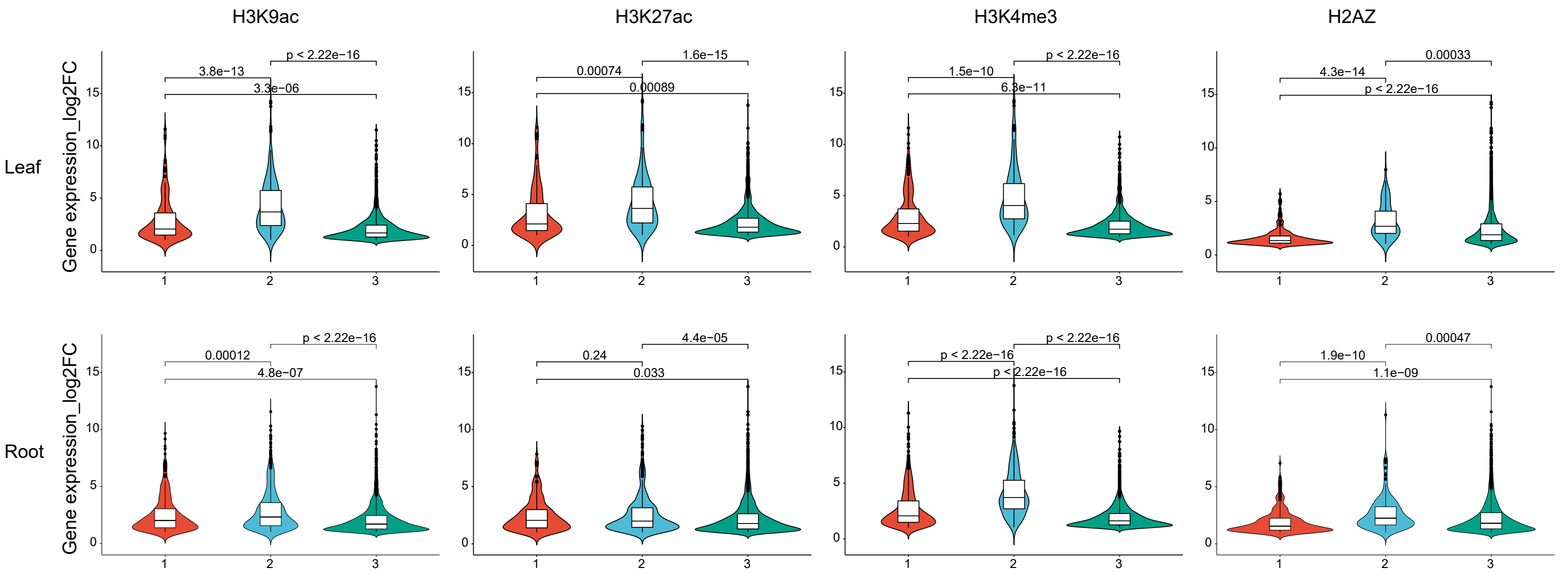


**Figure S8. The role of histone marks in the induction of PEG-responsive genes.**
1. Genes without histone mark enrichment;
2. Genes with increased (H3K9, H3K27, H3K4me3) or decreased (H2AZ) histone mark enrichent after PEG treatment;
3. Genes with unchanged histone mark enrichment after PEG treatment.

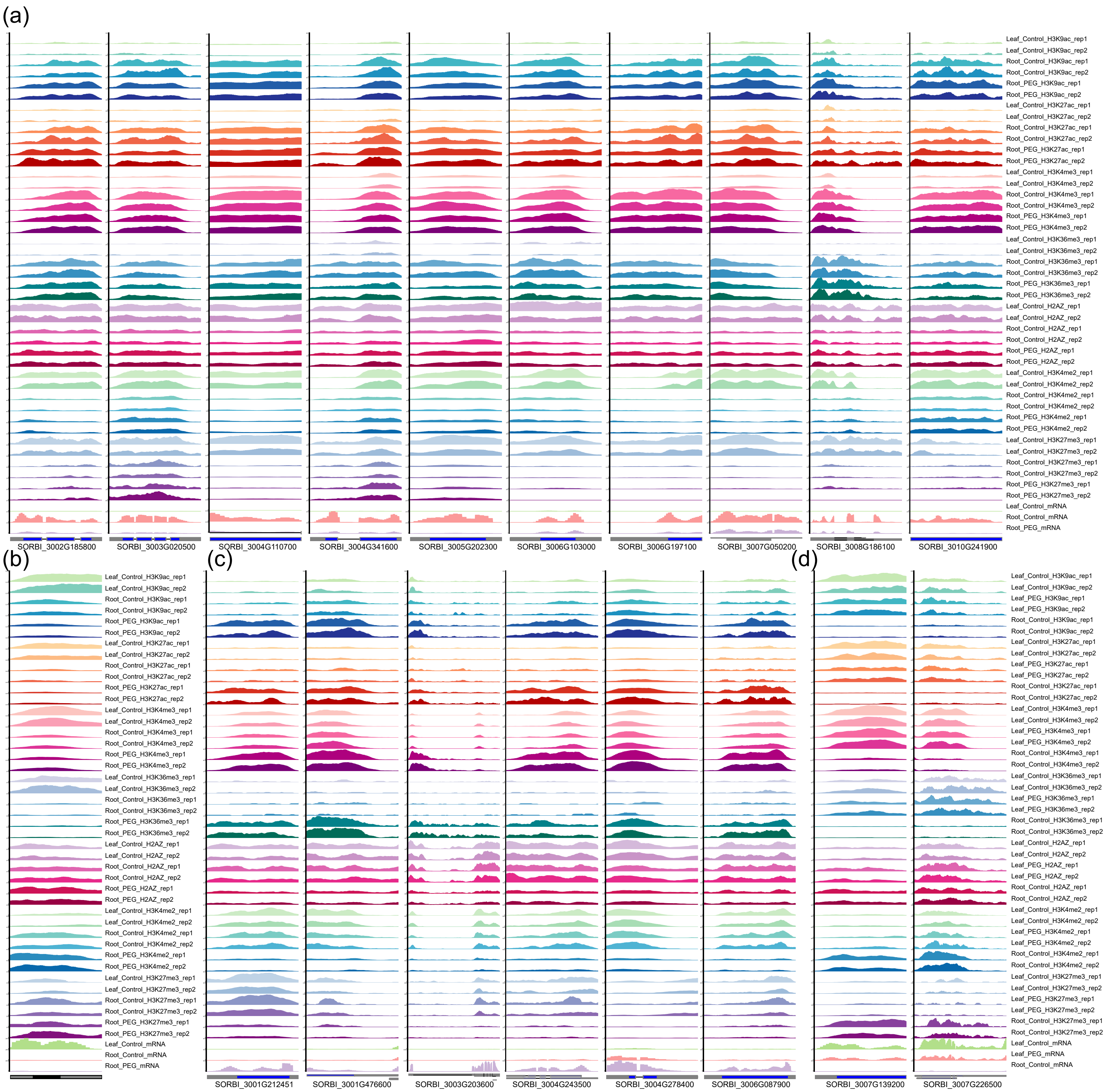


**Figure S9. Genome browser screen shots of histone mark enrichment in some common DEGs (identified in Figure 6) with 7 histone marks changing between leaf and root but not in response to PEG treatment.**
(**a**) Group IV; (**b**) Group III; (**c**) Group I; (**d**) Group II. For classification see Figure 7.

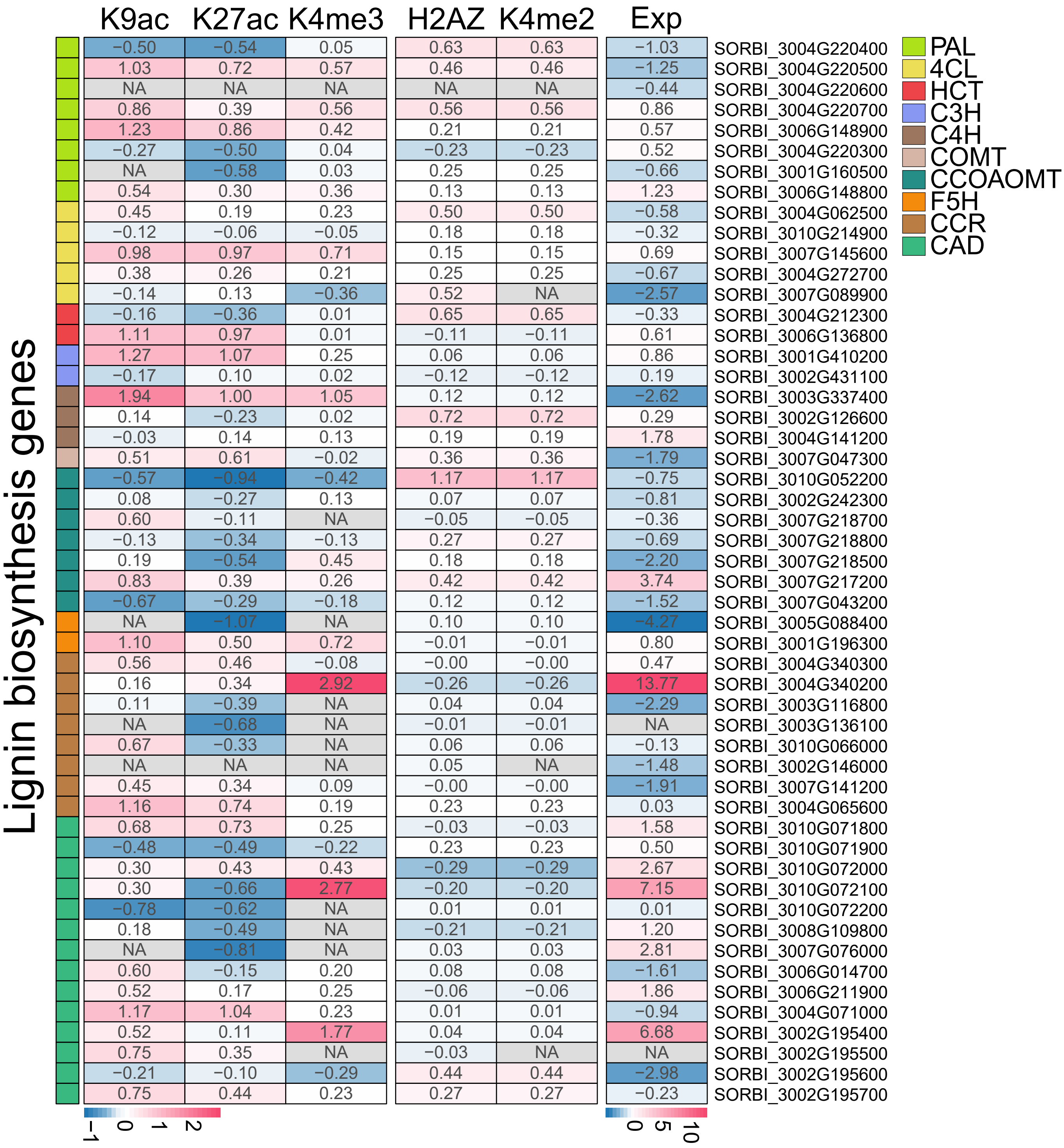


**Figure S10 Heat maps showing fold change (log2FC) of expression and histone marks of lignin biosynthesis genes**
